# Activity regulates a cell type-specific mitochondrial phenotype in zebrafish lateral line hair cells

**DOI:** 10.1101/2022.06.17.496604

**Authors:** Andrea McQuate, Sharmon Knecht, David W. Raible

## Abstract

Hair cells of the inner ear are particularly sensitive to changes in mitochondria, the subcellular organelles necessary for energy production in all eukaryotic cells. There are over thirty mitochondrial deafness genes, and mitochondria are implicated in hair cell death following noise exposure, aminoglycoside antibiotic exposure, as well as in age-related hearing loss. However, little is known about the basic aspects of hair cell mitochondrial biology. Using hair cells from the zebrafish lateral line as a model and serial block-face scanning electron microscopy, we have quantifiably characterized a unique hair cell mitochondrial phenotype that includes (1) a high mitochondrial volume, and (2) specific mitochondrial architecture: multiple small mitochondria apically, and a reticular mitochondrial network basally. This phenotype develops gradually over the lifetime of the hair cell. Disrupting this mitochondrial phenotype with a mutation in *opa1* impacts mitochondrial health and function. While hair cell activity is not required for the high mitochondrial volume, it shapes the mitochondrial architecture, with mechanotransduction necessary for all patterning, and synaptic transmission necessary for development of mitochondrial networks. These results demonstrate the high degree to which hair cells regulate their mitochondria for optimal physiology, and provide new insights into mitochondrial deafness.

## Introduction

Mitochondria are essential subcellular organelles in nearly all eukaryotic cells, where they perform and regulate manifold functions including ATP production, calcium buffering, apoptosis, metabolite generation, among others. These functions are influenced by a cell’s total mitochondrial volume, regulated by mitochondrial biogenesis (Jornayvaz and Shulman, 2010) and subsequent mitochondrial architecture, sculpted by mitochondrial fusion and fission (Picard et al., 2013). Mitochondrial fusion and elongated mitochondria are associated with heightened mitochondrial membrane potentials, increased ATP production, and improved calcium buffering (Picard et al., 2013; Szabadkai et al., 2006; Gomes et al., 2011). Meanwhile, mitochondrial fission and smaller mitochondria are associated with lower mitochondrial membrane potentials, lower ATP production, and apoptosis (Liu et al., 2020). The combination of these features produces an overall mitochondrial phenotype according to cellular need. Failure to achieve an appropriate mitochondrial phenotype results in a variety of pathologies, particularly in highly metabolically active cells such as electrically excitable cells (Reddy et al., 2011).

Hair cells in the peripheral auditory nervous system mediate hearing and balance. Deflection of the stereocilia bundle at the apical pole of the hair cell results in cation influx in a process known as mechanotransduction, and subsequent depolarization and calcium influx through voltage gated calcium channels results in glutamate release from ribbon synapses at the basolateral pole onto afferent neurons. Hair cells heavily depend on mitochondria to sustain energetic demands, with 75% of their ATP usage produced via oxidative phosphorylation (Puschner and Schacht, 1997) It is perhaps due to their high dependency on mitochondria that hair cells are particularly susceptible to mitochondrial alterations; mitochondria are implicated in both hereditary and environmentally induced hearing loss, as well as aging (Kokotas et al., 2007); Someya and Prolla, 2010; Böttger and Schacht, 2013).

Mitochondria are implicated at both poles of healthy hair cells. In rat cochlear inner hair cells, apical mitochondria have been shown to buffer calcium influx during mechanotransduction (Beurg et al., 2010; Pickett et al., 2018). Meanwhile, in zebrafish lateral line hair cells, basal mitochondrial calcium uptake is essential for regulating ribbon size (Wong et al., 2019). Mitochondria also play a role in hair cell vulnerability to aminoglycoside exposure. Hair cells that have been treated with neomycin demonstrate abnormal mitochondrial morphologies prior to other insults (Owens et al., 2007). Neomycin-induced hair cell death requires mitochondrial calcium uptake (Esterberg et al., 2014; 2016), and hair cell sensitivity to neomycin increases with cumulative mitochondrial activity (Pickett et al., 2018). Similarly, calcium import into the mitochondria via the MCU has been implicated in noise-induced hearing loss (Wang et al., 2019), and related mitochondrial potentials are disrupted in aging (Perkins et al., 2020) However, little is known regarding the morphological characteristics of hair cell mitochondria compared with other cell types, and how these morphologies intersect with hair cell function (Lesus et al., 2019). We hypothesized that hair cells maintained their mitochondria in an optimal configuration dependent on mitochondrial fusion to sustain high metabolic demands.

Here, we detailed the characteristics of mitochondria in the hair cells of the zebrafish lateral line. The lateral line is a powerful model system composed of clusters of hair cells called neuromasts. Lateral line hair cells are genetically and morphologically similar to the hair cells located within the mammalian inner ear, but are more easily accessible to genetic manipulations and experimentations (Pickett and Raible, 2019). As fluorescent markers lack the resolution to distinguish between individual organelles, we turned to serial block-face scanning electron microscopy (SBFSEM) to produce three-dimensional reconstructions of hair cells and their mitochondria with ultrastructural resolution. We demonstrated that zebrafish lateral line hair cells have a mitochondrial phenotype distinct from supporting cells. This phenotype includes a high total mitochondrial volume, and a particular architecture, with large, highly networked mitochondria at the basolateral pole near the synaptic ribbons, and smaller mitochondria positioned apically. This phenotype develops with hair cell maturation and is dependent on mechanotransduction and synaptic activity. Overall, our results demonstrate that hair cells might be sensitive to mitochondrial perturbations owing to the high specificity to which the organelle is regulated within the cell.

## Results

### Hair cells have a higher total mitochondrial volume than support cells

Although the importance of mitochondria in hair cells is well established, it is unclear how hair cell mitochondria compare with other cell types, both in number and morphology. We used SBFSEM to reconstruct hair cells (HC) in zebrafish neuromasts SO1 or SO2 (Raible and Kruse, 2000, Figure 1A). This technique provides sufficient resolution to distinguish between individual mitochondria and measure their volumes. It also allows comparison of individual mitochondrial morphologies. In our first set of experiments, we began serial sectioning about halfway through the neuromast (n = 3 neuromasts for HCs, n = 2 neuromasts for support cells, each neuromast from a different 5 dpf zebrafish). Hair cells (HC, Figure 1B) were easily distinguishable from other cell types by the presence of synaptic ribbons and stereocilia. Central support cells (Figure 1C) were defined as interdigitating between two hair cells, while peripheral cells (Figure 1D) could touch one or no hair cells. The roles of these two supporting cell types have been found to be different during homeostasis and regeneration, with peripheral cells contributing the bulk of progenitors fated to become hair cells, and central support cells serving a more “glial-like” function (Thomas and Raible, 2019; Romero-Carvajal et al., 2015). Mitochondria and cell bodies were reconstructed via manual segmentation. We found that hair cells contained 16.8 ± 1.1 µm^3^ of mitochondrial volume, distributed over 37 ± 1.7 individual mitochondria (Figure 1E and F). These measurements were significantly greater than in both central and peripheral support cells. Meanwhile, the average size of an individual mitochondrion only differed between hair cells and the peripheral supporting cells, with the peripheral cells having significantly smaller mitochondria than hair cells (Figure S1A). The ratio of mitochondria to cell volume was also significantly higher in hair cells than support cells (HCs, 9 ± 0.4% compared to central, 4 ± 0.3% or peripheral, 3 ± 0.2% SCs), eliminating the possibility that observed differences in mitochondrial volume were due to differences in cell size (Figure S1B).

**Figure 1:**
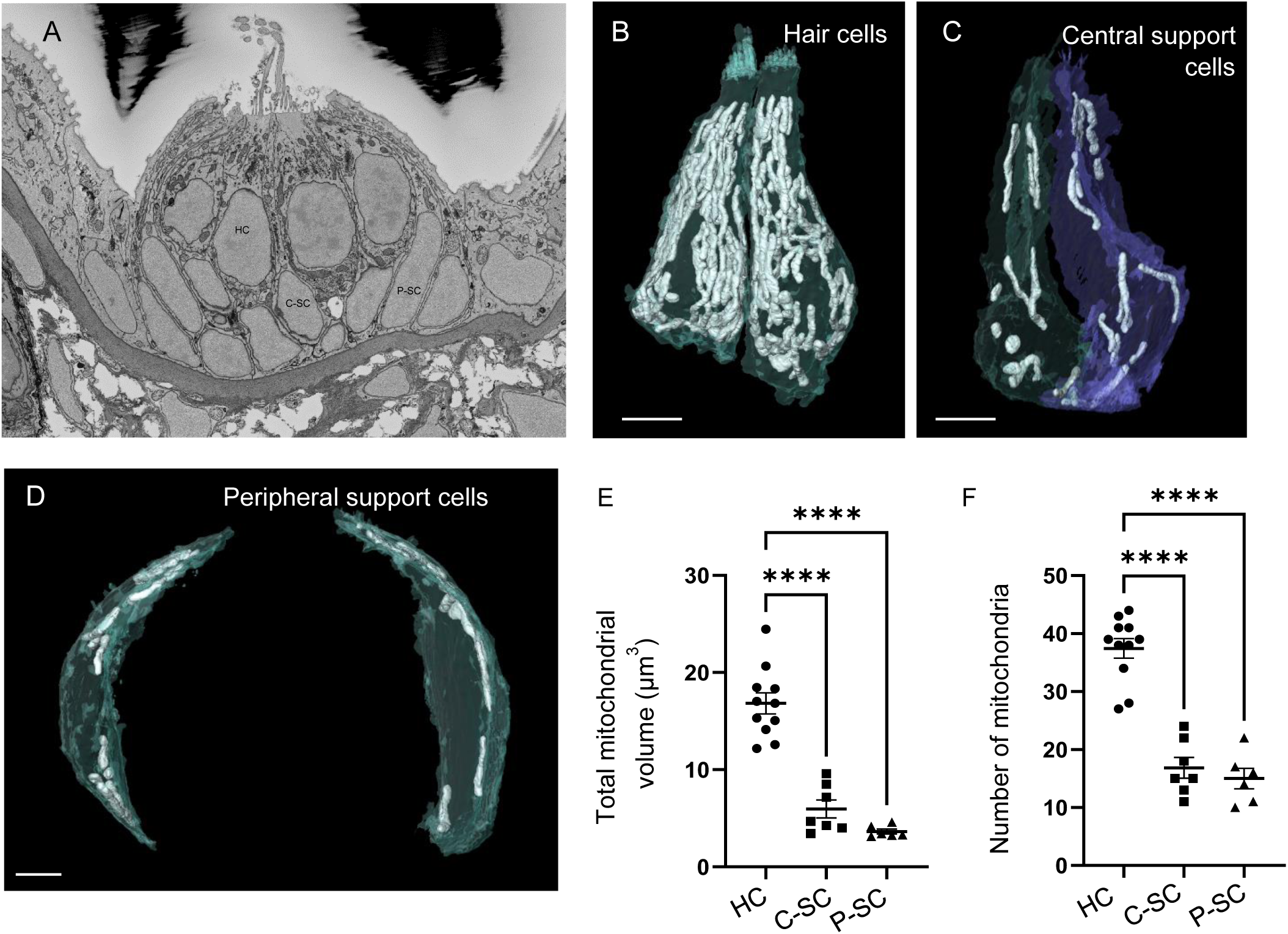
Hair cells contain a higher total mitochondrial volume than support cells. (A) SEM cross-section through 5dpf zebrafish NM SO1. (B) Reconstructed hair cells (HC) from NM SO1, with mitochondria shown in white. Scale bar = 3 µm. (C) Two central support cells (C-SC) reconstructed from NM SO1. Scale bar = 3 µm (D) Two peripheral support cells (P-SC) with mitochondria shown in white. Scale bar = 2 µm (E) Sum of mitochondrial volume for hair cells, central support cells, and peripheral hair cells. (In µm^3^) HC: 16.87 ± 1.1; C-SC: 5.9 ± 0.92; P-SC: 3.6 ± 0.24. (F) Number of individual mitochondria from hair cells, central support cells, and peripheral support cells. HC: 37.7 ± 1.7, C-SC: 16.9 ± 1.8, P-SC: 15 ± 1.8. For both (E) and (F) HC: n = 11, 3 NMs, 3 fish. C-SC: n = 7, 2 NMs, 2 fish. P-SC: n = 6, 1 NM, 1 fish. Data are presented as mean ± SEM; one-way ANOVA with Tukey’s multiple comparisons, **** p < 0.0001.

### Hair cell mitochondria have specific architecture

We found that hair cells did not contain a homogeneous mitochondrial population. Almost all hair cells contained a single interconnected mitochondrial network amid many other smaller mitochondria (Figure 2A). The mitochondrial networks (max mito) ranged from 2-10 µm^3^, on average 4-5 standard deviations from the mean mitochondrial volume (Figure 2B and C). In contrast, the bulk of hair cell mitochondria averaged ∼0.45 µm^3^. This made the largest mitochondria in each hair cell statistical outliers. Meanwhile, both central and peripheral support cells had homogeneous mitochondrial populations evenly distributed throughout the cells (Figure 2B and C). We found that hair cell mitochondrial networks preferentially localized to the basolateral compartment of hair cells, often surrounding the synaptic ribbons, though also extending apically (Figure 2A). On average, almost 70% of the mass of each large mitochondrion was located in the lowest quadrant of the hair cell (Figure 2D and E), consistent with active localization to the synaptic region. The remaining smaller mitochondria were positioned apically, and were more uniform in size, as shown by the linear nature of the cumulative distribution of mitochondrial volume from the apical cuticular plate to the basal ribbons (Figure 2F).

**Figure 2:**
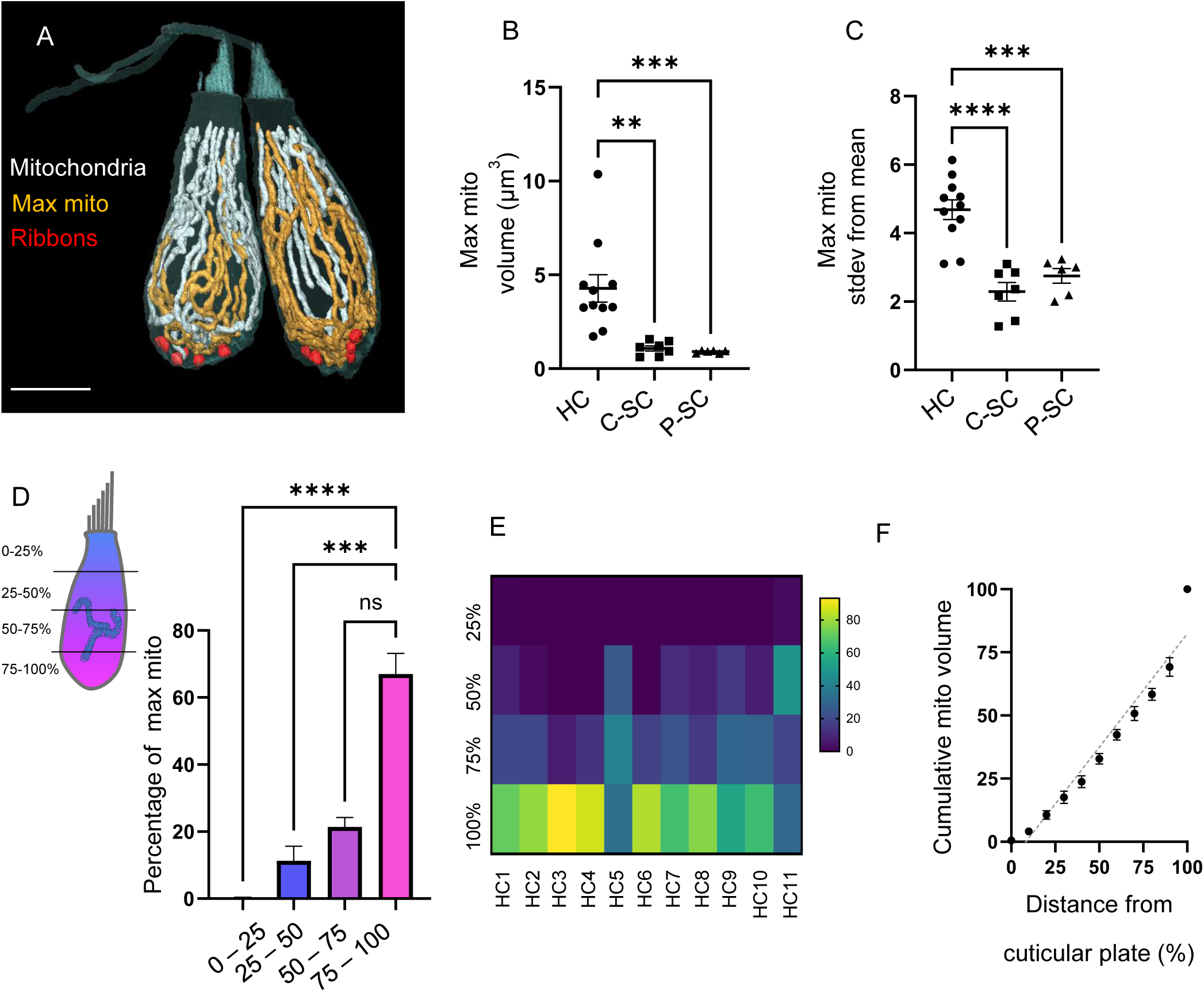
Hair cell mitochondria have a specific architecture. (A) Two reconstructed mature hair cells. Mitochondria are shown in white. Single largest mitochondrion shown in gold. Synaptic ribbons shown in red. Scale bar = 5 µm. (B) Volume of the largest mitochondrion in hair cells (HC), central support cells (C-SC), and peripheral support cells (P-SC). (In µm^3^) HC: 4.28 ± 0.7, C-SC: 1.1 ± 0.14, P-SC: 0.9 ± 0.03; mean ± SEM; Kruskal-Wallis test with Dunn’s multiple comparisons ** p = 0.003, *** p = 0.0009. (C) The number of standard deviations the largest mitochondrion is from the average, in hair cells, central support cells, and peripheral support cells. HC: 4.7 ± 0.28, C-SC: 2.3 ± 0.27. P-SC: 2.8 ± 0.21, mean ± SEM; one-way ANOVA with Tukey’s multiple comparison test, ****p < 0.0001, ***p = 0.0003. For (B - C) HC: n = 11, 3 NMs, 3 fish. C-SC: n = 7, 2 NMs, 2 fish. P-SC: n = 6, 1 NMs, 1 fish (Same cells as in Fig 1). (D) Summarized data for the position of the largest mitochondrion within hair cells. The length of each HC was normalized and broken into quadrants. The number of points the single largest mitochondrion was located within each cellular quadrant were counted and demonstrated as a percentage of the total. Summary data: 75-100: 67.01 ± 6.1%, 50-75: 21.4 ± 2.8%, 25-50: 11.3 ± 4.3%, 0-25: 0.24 ± 0.21, mean ± SEM. n = 11 HCs. Kruskal-Wallis test with Dunn’s multiple comparisons, ** p = 0.0052, **** p < 0.0001, *** p = 0.0006. (E) The same data shown in D represented as heat map for individual hair cells. (F) Cumulative distribution of mitochondrial volume as a function of position within the cell as measured as a percentage from the cuticular plate. Mitochondrial position was measured as its lowest point within the cell. Gray line: linear regression, R^2^ = 0.9, p < 0.0001.

### Hair cell mitochondrial architecture develops with maturation

In the next set of experiments, we reconstructed hair cells and mitochondria from whole neuromasts, beginning serial sectioning on the periphery of each neuromast from fish that were 3 dpf (n = 2 neuromasts, 4-6 HCs each), 5 dpf (n = 2 neuromasts, 10-12 HCs each), and 6 dpf (n = 2 neuromasts, 12-14 HCs each) (Figure 3A). We were then able to differentiate mitochondrial volume and architecture between young and mature hair cells. Ongoing, homeostatic turnover within zebrafish neuromasts guarantees that neuromasts will contain hair cells of different ages, with younger hair cells on the neuromast periphery, and mature hair cells in the center. We categorized hair cells as central or peripheral by calculating a z-score for the distance of the hair cells’ nucleus from the center of the neuromast; hair cells with nuclei > 1 z-score away from the center were considered peripheral, and those < 1 z-score away were considered central. The categorization of central vs. peripheral worked well for both 5 and 6 dpf neuromasts. As hair cells in 3 dpf neuromasts did not yet have a “concentric” patterning, and their hair cells were more “skewed” across the x-y axis, z-scores were first pooled across neuromasts to divide cells into more central or peripheral positions within the neuromasts. We found across all three ages, hair cells on the periphery of the neuromast contained significantly lower total mitochondrial volumes (Figure 3B, data pooled 3-6 dpf) and fewer mitochondria (Figure 3C) than hair cells in the center of the neuromast. The largest mitochondrion in peripheral hair cells was also smaller (Figure 3D) and closer to the mean (Figure 3E) than in central hair cells.

**Figure 3:**
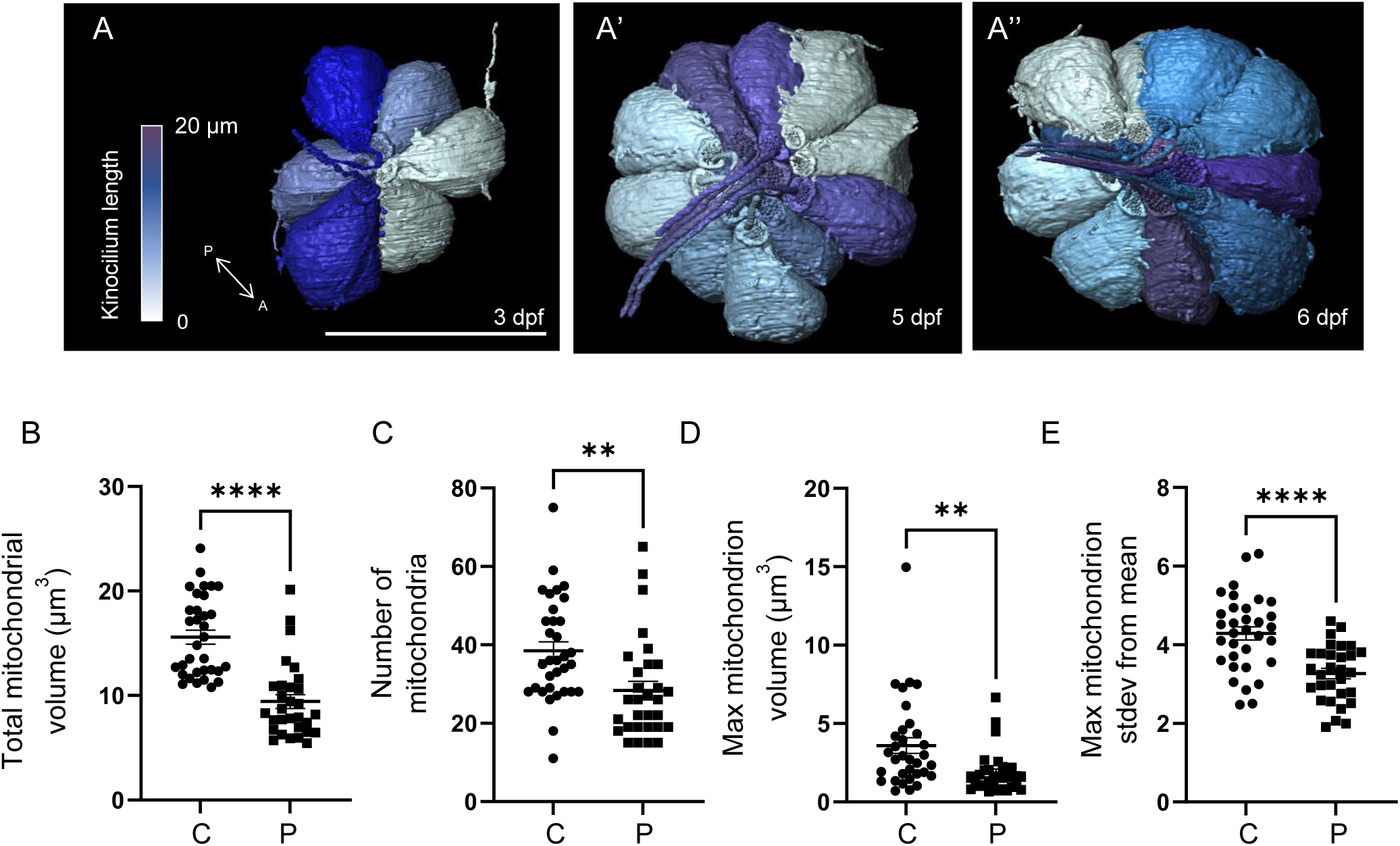
The hair cell mitochondrial phenotype develops during maturation. (A) Neuromast SO1 from 3 dpf (a), 5 dpf (a’), and 6 dpf (a’’) fish. Scale bar = 15 µm. Putative sister hair cell pairs are shown with matching colors according to kinocilium length. (B) Sum of total mitochondrial volumes in hair cells in the center of the neuromast (C) vs. on the periphery (P). (In µm^3^) Central: 15.59 ± 0.7, Peripheral: 9.4 ± 0.7. (C) Number of mitochondria in central vs peripheral hair cells. Central: 38.5 ± 2.3, Peripheral: 28.3 ± 2.4. (D) The volume of the largest mitochondrion in central vs. peripheral hair cells. (In µm^3^) Central: 3.6 ± 0.5, Peripheral: 1.8 ± 0.2. (E) The number of standard deviations the largest mitochondrion is from the average in central and peripheral hair cells. Central: 4.3 ± 0.2, Peripheral: 3.3 ± 0.13. For (B-E): Data include: 2, 3-dpf NMs (12 HCs total), 2, 5-dpf NMs (24 HCs total), 2, 6-dpf NMs (27 HCs total). Central HCs: n = 33. Peripheral HCs: n = 30. Presented as mean ± SEM. Kolmogorov-Smirnov test, **** p < 0.0001, ** p < 0.01.

As a continuous measure of hair cell development, we used the length of the kinocilium as a proxy for age, as documented previously (Kindt et al., 2012) in those blocks where full kinocilia remained after processing and serial sectioning (Figure 4A). We found a strong, positive correlation between the length of the kinocilium and the total mitochondrial volume (p < 0.0001, Figure 4B). We found that the number of individual mitochondria increased initially, but then plateaued as hair cells reached maturity, so that there was no significant correlation between kinocilium length and mitochondrial number (Figure 4C, p = 0.91). The growth of the large mitochondrial networks also correlated with kinocilium length (Figure 4D and E).

**Figure 4:**
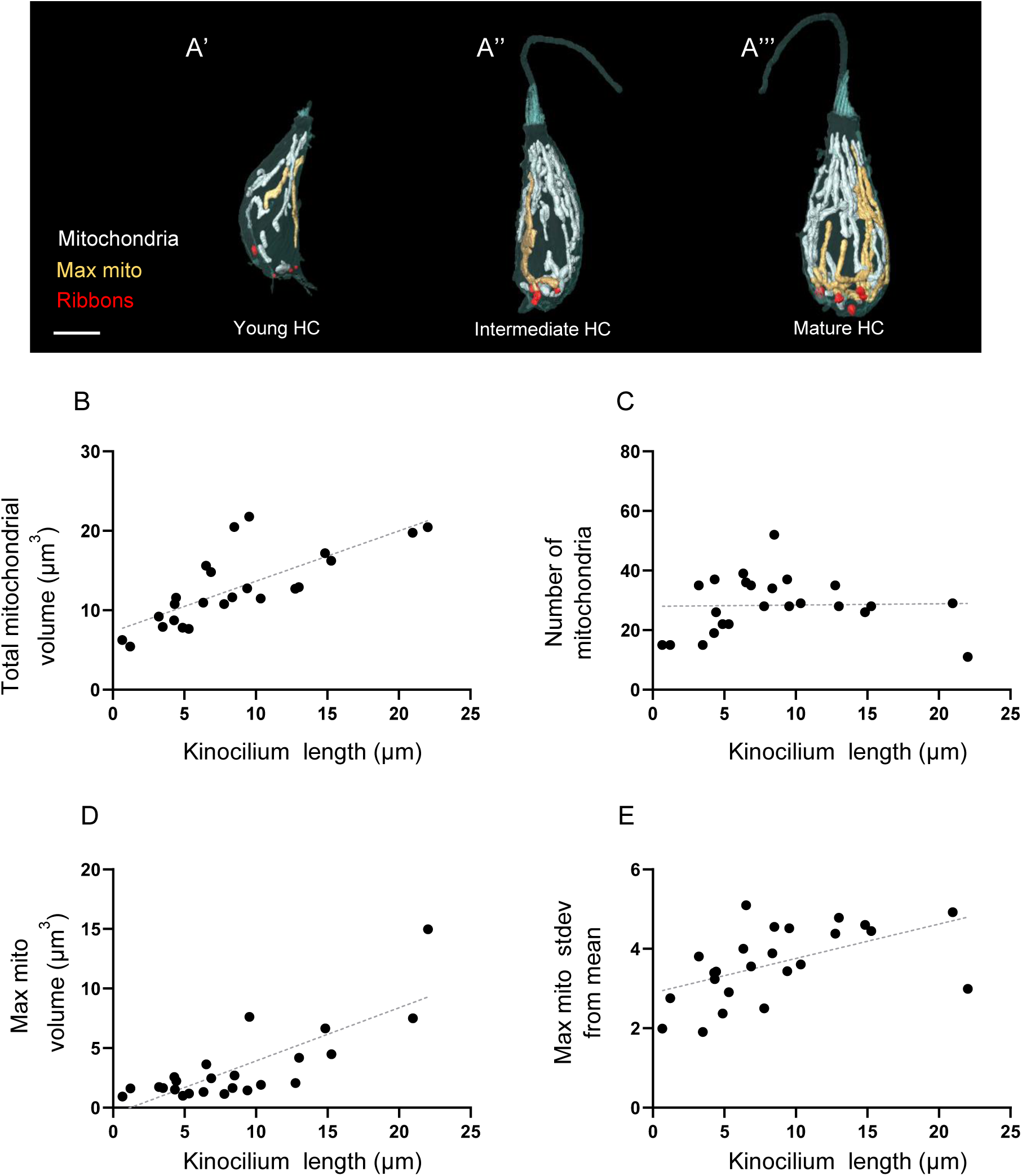
The hair cell mitochondrial phenotype correlates with kinocilium length. (A) Three hair cells of different ages from a 5 dpf WT neuromast. Mitochondria are shown in white. Single largest mitochondrion is shown in gold. Synaptic ribbons shown in red. Scale bar = 5 µm. (B) Relationship between kinocilium length and total mitochondrial volume. R^2^ = 0.57, p <0.0001. (C) Relationship between kinocilium length and the number of mitochondria. R^2^ = 0.0007, p = 0.9. (D) Relationship between kinocilium length and the volume of the largest mitochondrion. R^2^ = 0.6, p < 0.0001. (E) Relationship between kinocilium length and the max mito number of standard deviations from the mean. R^2^ = 0.27, p = 0.008. Data include 2, 5-dpf NMs from different fish, n = 24 HC total. For (B-E): Gray line = standard linear regression.

### Ribbon growth parallels mitochondrial growth and localization

Hair cell mitochondria are known to regulate ribbon volume, and their abilities to buffer calcium (Wong et al., 2019) and to generate ATP (Stowers et al., 2002; Perkins et al., 2010) at the basal end of hair cells plays roles in synaptic transmission. We reconstructed hair cell ribbons and analyzed their properties throughout hair cell maturation to better quantify their relationship with mitochondria. We found a steady increase in the total ribbon volume throughout hair cell development, both in the transition from peripheral to central hair cells, and across multiple ages of fish (Figure 5A). This was not attributable to a change in the number of ribbons (∼5, Figure 5B), which remained relatively stable, but due to an increase in the average ribbon size (Figure 5C). Plotting ribbon properties against kinocilium length yielded a similar result (Figure 5D-F). The average ribbon volume per hair cell climbed in a non-linear fashion before reaching a plateau (R = 0.7, Figure 5F). These changes in the hair cell ribbons mirrored changes in the mitochondria. The total mitochondrial volume correlated with total ribbon volume (Figure S2A). The largest mitochondrion also localized to the ribbons, as reflected in a non-linear decrease in the average minimum distance between the single largest mitochondrion and the ribbons (See Methods, Figure 5G, p = 0.001). Across all three ages of fish, the largest hair cell mitochondrion localized closer to ribbons than the median mitochondrion. However, the difference between the largest mitochondrion and the median mitochondrion grew larger with development (Figure 5H). At 3 dpf, there was an average difference of 3.4 µm. In peripheral immature hair cells of neuromasts of either 5 or 6 dpf fish, that difference was 2.2 µm. In central mature hair cells from 5 or 6 dpf fish, the difference grew to 6.2 µm. This suggests the active localization of the largest mitochondrion to the ribbons over the course of maturation.

**Figure 5:**
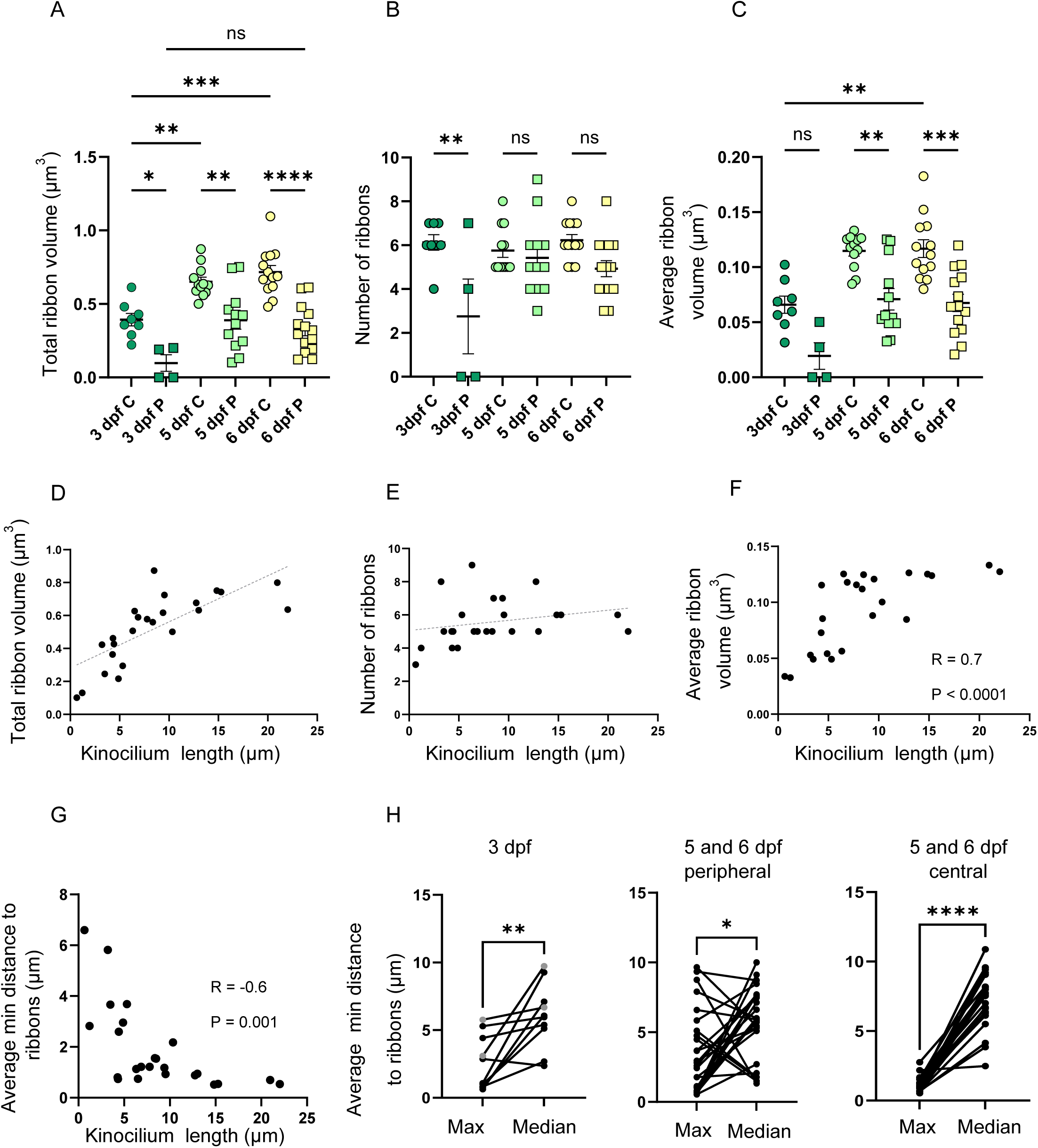
The largest mitochondrion localizes to the ribbons during hair cell maturation. (A) The total ribbon volume per central vs. peripheral hair cells across multiple ages. (In µm^3^) 3 dpf central: 0.39 ± 0.04, 3 dpf peripheral: 0.1 ± 0.06, 5 dpf central: 0.65 ± 0.03, 5 dpf peripheral: 0.39 ± 0.06, 6 dpf central: 0.72 ± 0.04, 6 dpf peripheral: 0.3 ± 0.04. (B) Number of ribbons per hair cell. 3 dpf central: 6.1 ± 0.4, 3 dpf peripheral: 2.7 ± 1.7, 5 dpf central: 5.7 ± 0.3, 5 dpf peripheral: 5.4 ± 0.5, 6 dpf central: 6.2 ± 0.3, 6 dpf peripheral: 4.9 ± 0.4. (C) Average ribbon volume per central vs. peripheral hair cell across multiple ages. (In µm^3^) 3 dpf central: 0.06 ± 0.008, 3 dpf peripheral: 0.02 ± 0.01, 5 dpf central: 0.11 ± 0.005, 5 dpf peripheral: 0.07 ± 0.01, 6 dpf central: 0.12 ± 0.008, 6 dpf peripheral: 0.07 ± 0.008. For (A-C), data are mean ± SEM, one-way ANOVA with Tukey’s multiple comparisons, * p < 0.05, ** p < 0.01, *** p < 0.005, **** p < 0.0001. 3 dpf central, n = 8 HCs, 3 dpf peripheral, n = 4 (2 neuromasts from different fish) 5 dpf central, n = 12, 5 dpf peripheral, n = 12 (2 neuromasts from different fish), 6 dpf central n = 13, 6 dpf peripheral n = 14 (2 different neuromasts from 1 fish). (D) Relationship between kinocilium length and total ribbon volume. R^2^ = 0.55, p < 0.0001. (E) Relationship between kinocilium length and the number of ribbons. R^2^ = 0.07, p = 0.2. (D and E) Gray line = linear regression. (F) Relationship between kinocilium length and average ribbon volume per hair cell, Pearson’s correlation. (D – F) n = 24 HCs (2, 5 dpf neuromasts from two different fish). (G) Relationship between kinocilium length and average minimum distance to the largest mitochondrion. Pearson’s correlation. Data include 2, 5 dpf NMs from different fish, n = 24 HC total. (H, Left) Average minimum distance between the max or median mitochondrion and the synaptic ribbons (center to center) in 3 dpf animals, for both central and peripheral hair cells. Gray points denote peripheral hair cells, black points denote central hair cells. In one neuromast, the pair of peripheral hair cells contained no ribbons, precluding this analysis. (In µm) Max: 2.5 ± 0.6, median 6 ± 0.8, mean ± SEM, n = 10 HCs, 2 NMs (same as in Fig 3), paired t-test **p = 0.006. (H, middle and right) Average minimum distance between max or median mitochondrion and the synaptic ribbons in 5-6 dpf peripheral HCs (middle) or central hair cells (right). Max peripheral: 3.5 ± 0.6, median peripheral: 5.8 ± 0.5, mean ± SEM, n = 26 HCs (2, 5dpf NMs with 12 HCs; 2, 6dpf NMs with 14 HCs), paired t-test *p = 0.01. Max central: 1.1 ± 0.1, median central: 7.3 ± 0.38, mean ± SEM, n = 25 HCs, (2, 5dpf NMs with 12 HCs; 2, 6dpf NMs with 13 HCs), paired t-test **** p < 0.0001.

### Multidimensional analysis of hair cell mitochondrial and ribbon properties shows trajectory of mitochondrial development

To quantifiably assess the maturation of the hair cell mitochondrial phenotype, we performed a principal component analysis regarding seven aspects of hair cell mitochondria and synaptic ribbons: (1) number of mitochondria, (2) the total mitochondrial volume, (3) volume of the largest mitochondrion, (4) volume of the median mitochondrion, (5) average minimum distance of largest mitochondrion to the ribbons, (6) average minimum distance of the median mitochondrion to the ribbons, (7) the total ribbon volume, and (8) average ribbon volume. Using the first six principal components which described roughly 95% of the variability in the data set, we then plotted the results in a two-dimensional UMAP space. Hair cells were color-coded according to the length of their kinocilium. Results showed the gradual transition between young and mature hair cell mitochondrial properties (Figure 6A), with kinocilium length showing spatial correlation across the manifold (Moran’s I = 0.5, p = 0.001). We also observed a separation of mitochondrial properties across the three ages (Figure 6B). These results quantifiably demonstrate the trajectory of hair cell mitochondrial maturation, consistent with a gradual development of this phenotype.

**Figure 6:**
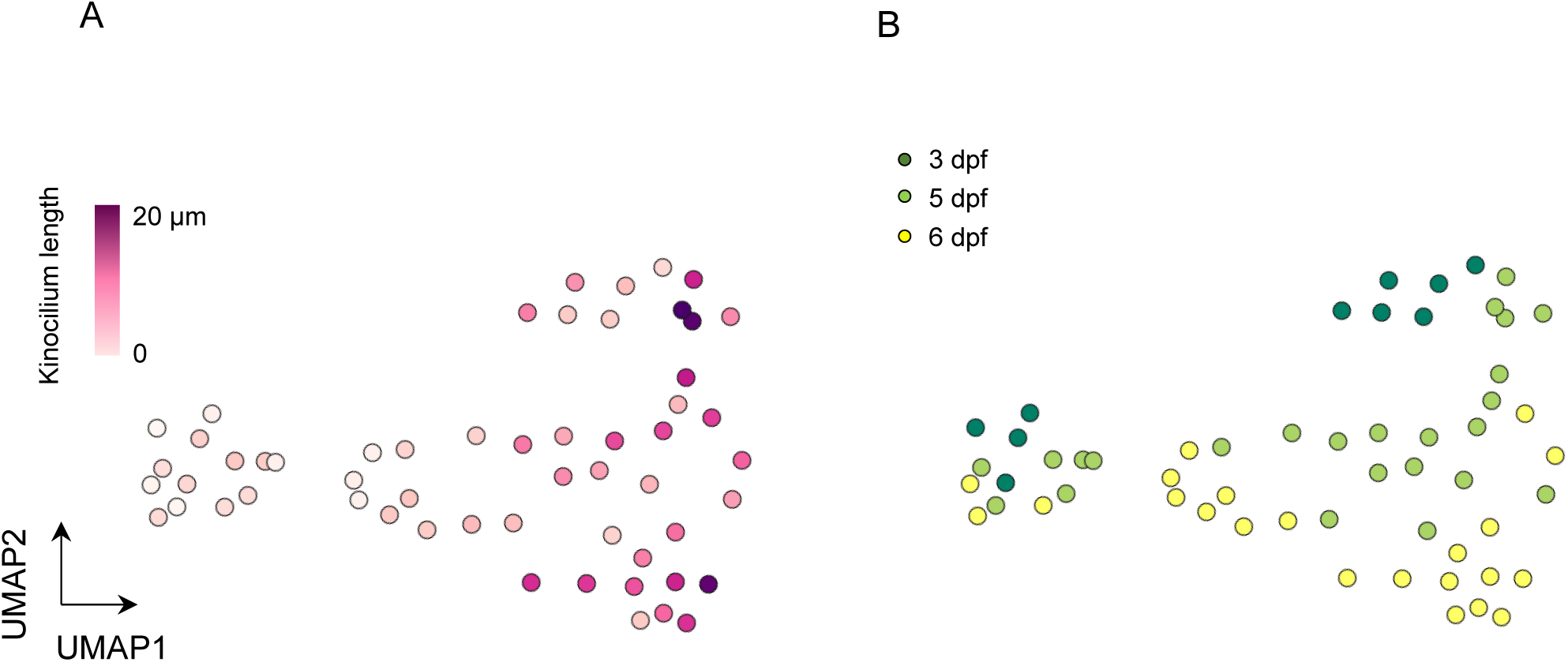
Hair cell mitochondrial phenotype can be quantifiably identified. (A) UMAP plot of hair cells based on principal component analysis of mitochondrial properties. Variables included in this analysis: (1) Number of mitochondria, (2) Total mitochondrial volume, (3) Max mito volume, (4) Median mito volume, (5) Average minimum distance between max mito and ribbons, (6) Average minimum distance between median mito and ribbons, (7) Total ribbon volume, (8) Average ribbon volume. Hair cells are color-coded by the length of their kinocilium. (B) Same UMAP analysis as in A, color-coded according to age of the animal. Total HCs, n = 56. 3-dpf HC: n = 10, 5-dpf HC: n = 24, 6-dpf HC: n = 22.

### Disrupting hair cell mitochondrial architecture impacts their function

To test the impact of mitochondrial architecture on hair cell function, we created a CRISPR mutant for the gene *opa1,* a dynamin-like GTPase necessary for the fusion of the inner mitochondrial membrane. The mutant was generated by introducing a premature stop codon into the second exon, resulting in a truncated, non-functional protein. Hair cells of these mutants completely lacked the mitochondrial architecture found in WT (Figure 7A), with a four-fold greater number of individual mitochondria (156 ± 23) and no mitochondrial network (Figure 7B). The total mitochondrial volume, however, was unaffected (Figure 7C). As smaller mitochondria are associated with lower mitochondrial membrane potentials and decreased OXPHOS capacity, we used TMRE dye to measure mitochondrial membrane potential in *opa1* mutants (Figure 7D). *Opa1* mutant hair cell mitochondria took up less of the TMRE dye than WT (Figure 7E), reflecting depolarized and less effective mitochondria. We also looked at mitochondrial calcium uptake during waterjet stimulation in *opa1* hair cells. Fish were transgenic for GCaMP3 targeted to the inner mitochondrial matrix under a hair cell-specific promoter (*Tg(myo6:mitoGCaMP3),* Figure 7F). A sinusoidal pressure wave of 10 Hz was applied to the stereocilia bundle (see Pickett et al., 2018). Larvae were genotyped following calcium imaging. Mitochondria from *opa1* hair cells had on average reduced GCaMP responses to waterjet, indicating that they took up less calcium than siblings (Figure 7G and H). These results suggest that the development of networked mitochondria through mitochondrial fusion assist with calcium uptake during mechanotransduction.

**Figure 7:**
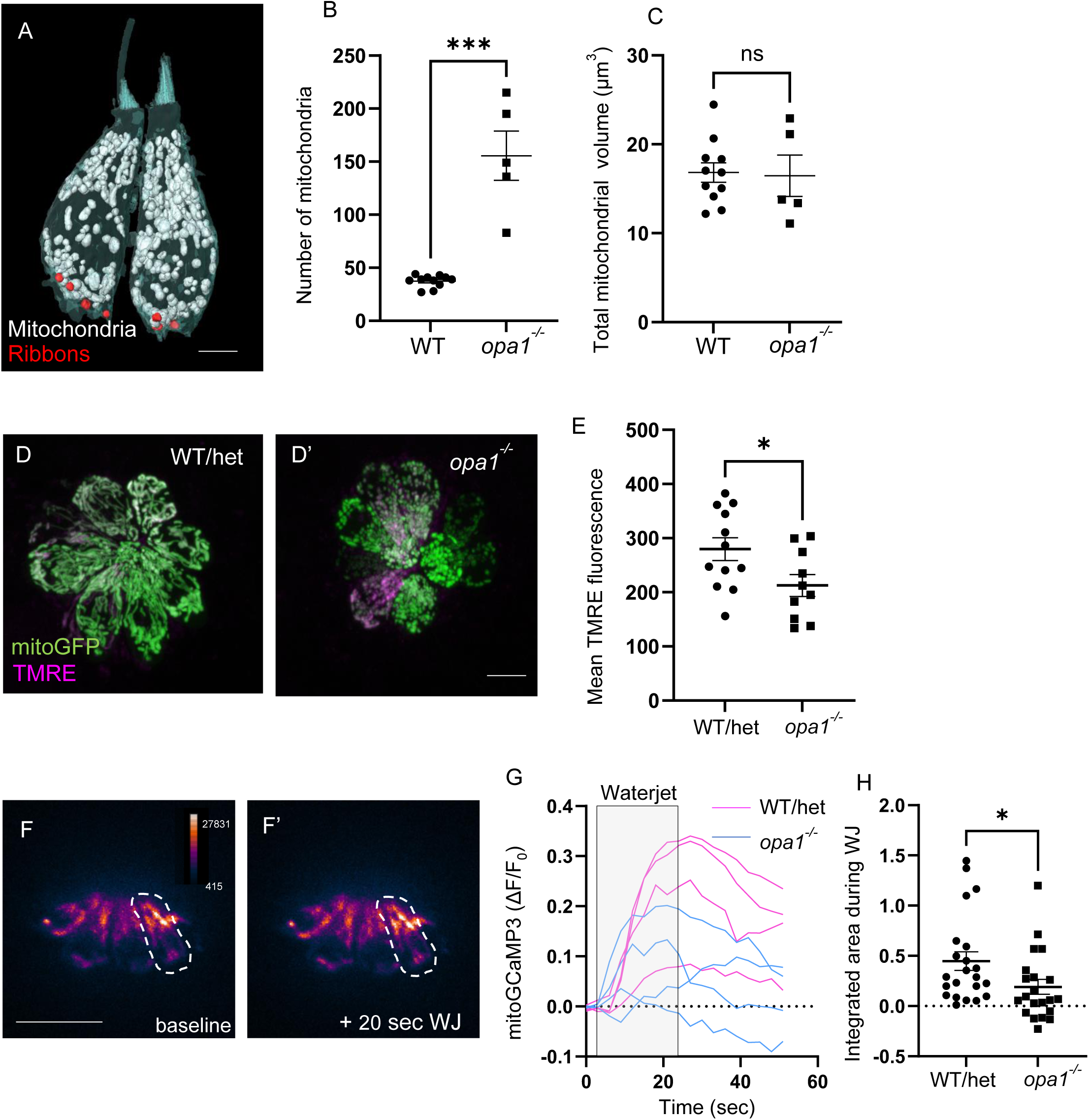
Mutations in *opa1* disrupt hair cell mitochondrial architecture and function. (A) Two reconstructed *opa1* hair cells. Mitochondria are shown in white. Synaptic ribbons in red. Scale bar = 5 µm. (B) Number of individual mitochondria in WT hair cells vs *opa1* hair cells. WT: 37.4 ± 1.7, *opa1*: 155.6 ± 23, mean ± SEM. Kolmogorov-Smirnov test, *** p < 0.001. (C) Sum of the mitochondrial volume in WT vs *opa1* hair cells. (In µm^3^) WT: 16.8 ± 1.1, *opa1*: 16.5 ± 2.3, mean ± SEM. Unpaired t-test, p = 0.87. For both (B-C), WT HC, n = 11 (3 NMs, 3 fish, same as Figures 1 and 2), *opa1* HC n = 5 (1 NM, 1 fish). (D) Confocal images of whole neuromasts containing *myo6b:mitoGFP* and incubated in 1 nM TMRE for 1 hr. The GFP signal was used to render a 3D mask, by which TMRE florescence was measured. Scale bar = 5 µm. (E) Average TMRE fluorescence in whole WT and *opa1* neuromasts. (In AU) WT: 279.6 ± 21.2, *opa1*: 212.6 ± 20.24, mean ± SEM. WT n = 12 NMs (5 fish), *opa1* n = 10 NMs (3 fish). Unpaired t-test, * p < 0.05. (F) WT hair cells expressing Tg(*myo6b:mitoGCaMP3*) indicator at baseline and at the end of a 20 sec, 10 Hz waterjet. Dotted line indicates example ROI of a single hair cell responding to waterjet stimulus. Scale bar = 10 µm. (G) Example mean myo6b:mitoGCaMP3 signal (expressed as ΔF/F_0_) following a 20 sec, 10 Hz waterjet. (H) The integrated charge transfer for WT/het and *opa1* hair cell mitochondria during the time of WJ stimulus. WT/het: (In AU) 0.4 ± 0.09, *opa1*: 0.2 ± 0.07, mean ± SEM. WT/het HC, n = 22 cells, 9 fish. *opa1* HC = 21 cells, 7 fish. Unpaired t-test, * p < 0.05.

### Hair cell activity regulates the development of mitochondria architecture

In many cell types, including skeletal muscle and cardiomyocytes, activity and intracellular calcium drive mitochondrial biogenesis to support an upregulated metabolic load (Ojuka et al., 2002; Chin, 2004). We hypothesized that development of mechanotransduction activity during hair cell maturation resulted in an increased energetic load that would similarly drive hair cell mitochondrial biogenesis and patterning. We therefore asked whether mechanotransduction was necessary for either the high mitochondrial volume or the specific mitochondrial architecture within hair cells. We reconstructed neuromasts from a 5 dpf *cdh23* mutant zebrafish (Figure 8A, n = 2 neuromasts), which lacks the tip-links necessary to open mechanotransduction channels (Söllner et al., 2004). The kinocilia of these hair cells were lost upon serial sectioning, hindering a full developmental analysis. In addition, hair cells failed form a pattern of central and peripheral hair cells, with a skewed arrangement more similar to the 3 dpf neuromasts. Therefore, we used the WT half-neuromast data set in this comparison (Figures 1-2). Surprisingly, *cdh23* mutant hair cells had similar total mitochondrial volumes as WT hair cells, implying that mechanotransduction was not necessary to drive the high mitochondrial volumes in these cells (Figure 8B). However, *cdh23* hair cell mitochondria were on average larger and more uniform than WT hair cells, with reduced mitochondrial number (Figure 8C). Although there was no significant difference in volume between the largest mitochondrion in *cdh23* and WT hair cells, the largest mitochondria in *cdh23* hair cells were not outliers, as the population of mitochondria was more uniformly large (Figure 8D). The relationship between the total mitochondrial volume and total ribbon volume was also disrupted in *cdh23* hair cells (p = 0.9, Figure S2B). *cdh23* hair cells contained similar ribbon numbers as WT (5.0 ± 0.3 ribbons). Ribbons were slightly smaller (0.08 ± 0.007 µm^3^ compared to 0.11 ± 0.007 in WT, p = 0.002), resulting in a smaller total ribbon volume (0.38 ± 0.03 compared to 0.54 ± 0.04 in WT, p = 0.005). The largest mitochondrion often spanned the entire cell, so there was no difference in its minimum distance of the largest mitochondrion to ribbons. However, unlike in WT, the largest mitochondrion in *cdh23* mutants showed no preference for the bottom quadrant of the hair cell (Figure 8F and G), with a significantly lower percentage located in the basolateral quadrant than WT (p < 0.001, two-way ANOVA). This implies that while mechanotransduction is not necessary for the high hair cell mitochondrial volume, it is necessary for the mitochondrial patterning, including an increase in the number of smaller mitochondria and the positioning of the largest mitochondrion preferentially near the ribbons.

**Figure 8:**
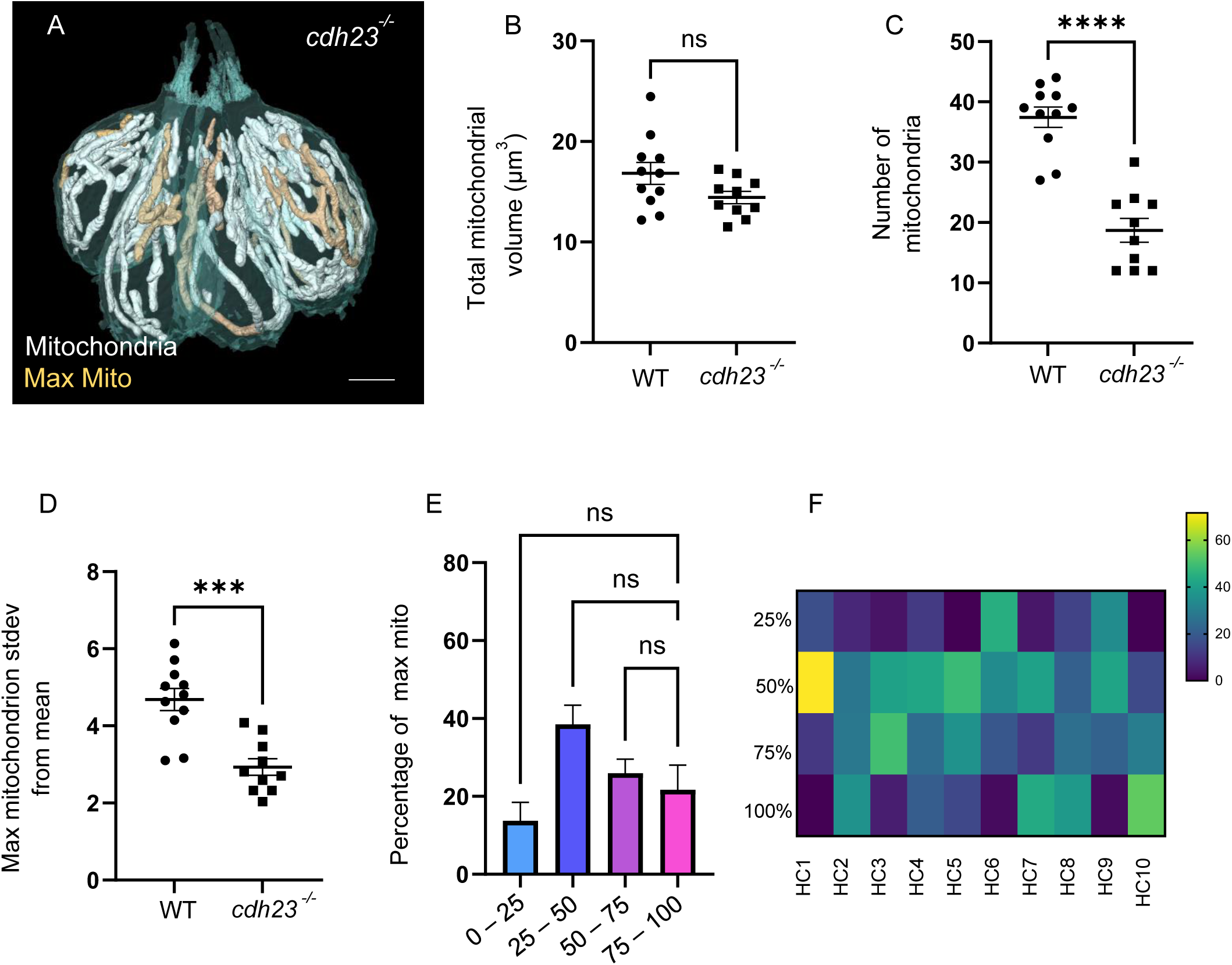
Mechanotransduction is not necessary for high mitochondrial volume, but all mitochondrial architecture. (A) Reconstructed HCs from a *cdh23* neuromast. Mitochondria shown in white. Single largest mitochondrion shown in gold. Scale bar = 5 µm. (B) Total mitochondrial volumes in WT and *cdh23* hair cells. (In µm^3^) WT: 16 ± 1.1, *cdh23*: 14.43 ± 0.6, mean ± SEM. Unpaired t-test, p = 0.07. (C) Number of mitochondria in WT and *cdh23* hair cells. WT: 37.45 ± 1.7, *cdh23*: 18.7 ± 2.0, mean ± SEM. Unpaired t-test, ****p < 0.0001. (D) Number of standard deviations the largest mitochondrion lies from the average volume in WT and *cdh23* hair cells. WT: 4.7 ± 0.3, *cdh23*: 3 ± 0.2, mean ± SEM. Unpaired t-test, ***p = 0.0001. (For B – D): WT, n = 11 (3 NMs, 3 fish, same as in Figures 1 and 2), *cdh23*, n = 10 (2 NMs, 1 fish). (E) Summary data of the percentage of the largest *cdh23* mitochondrion located within each cellular quadrant. 100-75%: 21.73 ± 6.3, 75-50%: 26.0 ± 3.6, 50-25%: 38.5 ± 5.0, 25-0%: 13.8 ± 4.7, mean ± SEM. n = 10 HCs. Kruskal-Wallis test with Dunn’s multiple comparisons, non-significant. (F) Individual cell data showing the percentage of the largest mitochondrion located within each hair cell quadrant.

The consistent localization of the largest mitochondrion to the ribbons implies a role in synaptic transmission. We therefore next asked whether synaptic transmission was necessary for the growth of the mitochondrial network. To this end, we reconstructed hair cells from a 5 dpf *cav1.3a* mutant (Figure 9A, n = 2 neuromasts), which lack the L-type calcium channel necessary for the calcium influx resulting in neurotransmitter release (Trapani and Nicolson, 2011). The full kinocilia of these hair cells was preserved upon serial sectioning, allowing analysis of their developmental trajectory. *cav1.3a* hair cells developed mitochondrial volume with age (p < 0.0001, Figure 9B), though the growth in mitochondrial volume was significantly faster than in WT (difference in slopes, p < 0.05). However, the largest mitochondrion in *cav1.3a* mutant hair cells did not expand over development (Figure 9D, p = 0.6, difference from WT, p < 0.0001). This paralleled an unrestrained rise in the number of individual mitochondria (Figure 9E, p < 0.0001, difference from WT, p < 0.0001). Furthermore, although the largest mitochondrion in *cav1.3a* mutant hair cells showed a preference for the most basolateral quadrant similar to WT (not shown), there was no significant localization of the largest mitochondrion to ribbons (Figure 9F, p = 0.08), which may correspond to the reductions in the size of this mitochondrion. These data imply that calcium influx through the CaV1.3 channel might drive growth and localization of the largest mitochondrial network. *cav1.3a* hair cells across development had significantly more ribbons (7 ± 0.3, compared to 5.6 ± 0.4 in WT, p = 0.002), but there was no difference in the total ribbon volume. Instead, in hair cells that were mature (kinocilia longer than 5 µm, see Figure 5), we found *cav1.3a* cells consistently had slightly smaller ribbons on average (WT, 0.11 ± 0.006 µm^3^, *cav1.3a*, 0.09 ± 0.004 µm^3^, p = 0.01). The *cav1.3a* ribbons could be found in tightly packed clusters (Figure S3B). Despite being smaller on average, *cav1.3a* cells also contained some large ribbons with irregular patterning (Figure S3B). However, the relationship between total mitochondrial volume and total ribbon volume remained (p < 0.0001, Figure S2C).

**Figure 9:**
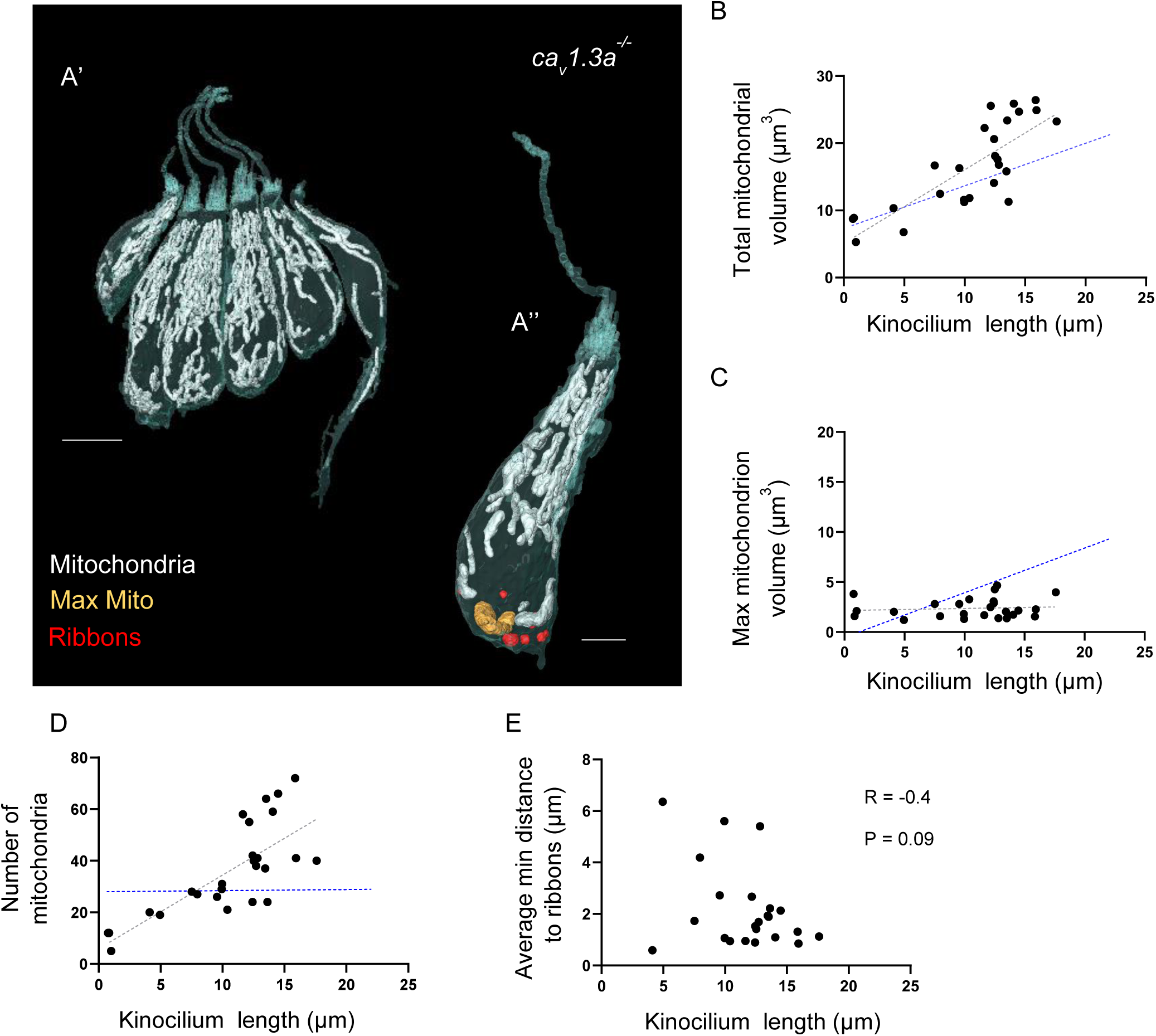
Synaptic transmission is necessary for growth of the largest mitochondrial network. (A) (a’) Six hair cells reconstructed from a *cav1.3a* neuromast, with mitochondria shown in white. Scale bar = 5 µm. (a’’) Single *cav1.3a* hair cell, mitochondria shown in white, single largest mitochondrion shown in gold, synaptic ribbons shown in red. Scale bar = 3 µm. (B) Relationship between *cav1.3a* kinocilium length and total mitochondrial volume. Gray line: linear regression: R^2^ = 0.6, p < 0.0001. Blue line: WT regression, difference in slope: p < 0.05. (C) Relationship between *cav1.3a* kinocilium length and volume of the max mitochondrion *cav1.3a* hair cells. Gray line: linear regression, R^2^ = 0.009, p = 0.6. Blue line: WT regression, difference in slopes: p < 0.0001. (D) Relationship between *cav1.3a* kinocilium length and the number of mitochondria. Gray line: linear regression, R^2^ = 0.6, p < 0.0001. Blue line: WT regression: difference in slopes: p < 0.0001. (E) Relationship between *cav1.3a* kinocilium length and the average minimum distance between the largest mitochondrion and the synaptic ribbons. Pearson’s correlation: R = -0.4, p =0.09. For (B – D): n = 25 HCs. For (F): n = 23 (two HCs did not yet contain ribbons). All *cav1.3a* cells are from 2, 5 dpf NMs, 1 fish. WT data are the same as in Figure 4.

To evaluate how both mechanotransduction and synaptic activity regulate the development of the hair cell phenotype, we included both mutants in the principal component analysis (Figure 10A, all genotypes combined). In Figures 10B and C, WT data were compared to either *cdh23* or *cav1.3a* data, respectively. Nearest neighbor analysis demonstrated that both *cdh23* and *cav1.3a* mutant hair cells were more likely to cluster with themselves (p = 0.001, both *cdh23* and *cav1.3a*) than with WT (p = 1.0 both *cdh23* and *cav1.3a*). In contrast, hair cells from *cdh23* siblings were not statistically likely to cluster apart from WT (p = 0.4). These data demonstrate that mutations in either mechanotransduction or synaptic transmission disrupt proper development of the hair mitochondrial phenotype.

**Figure 10:**
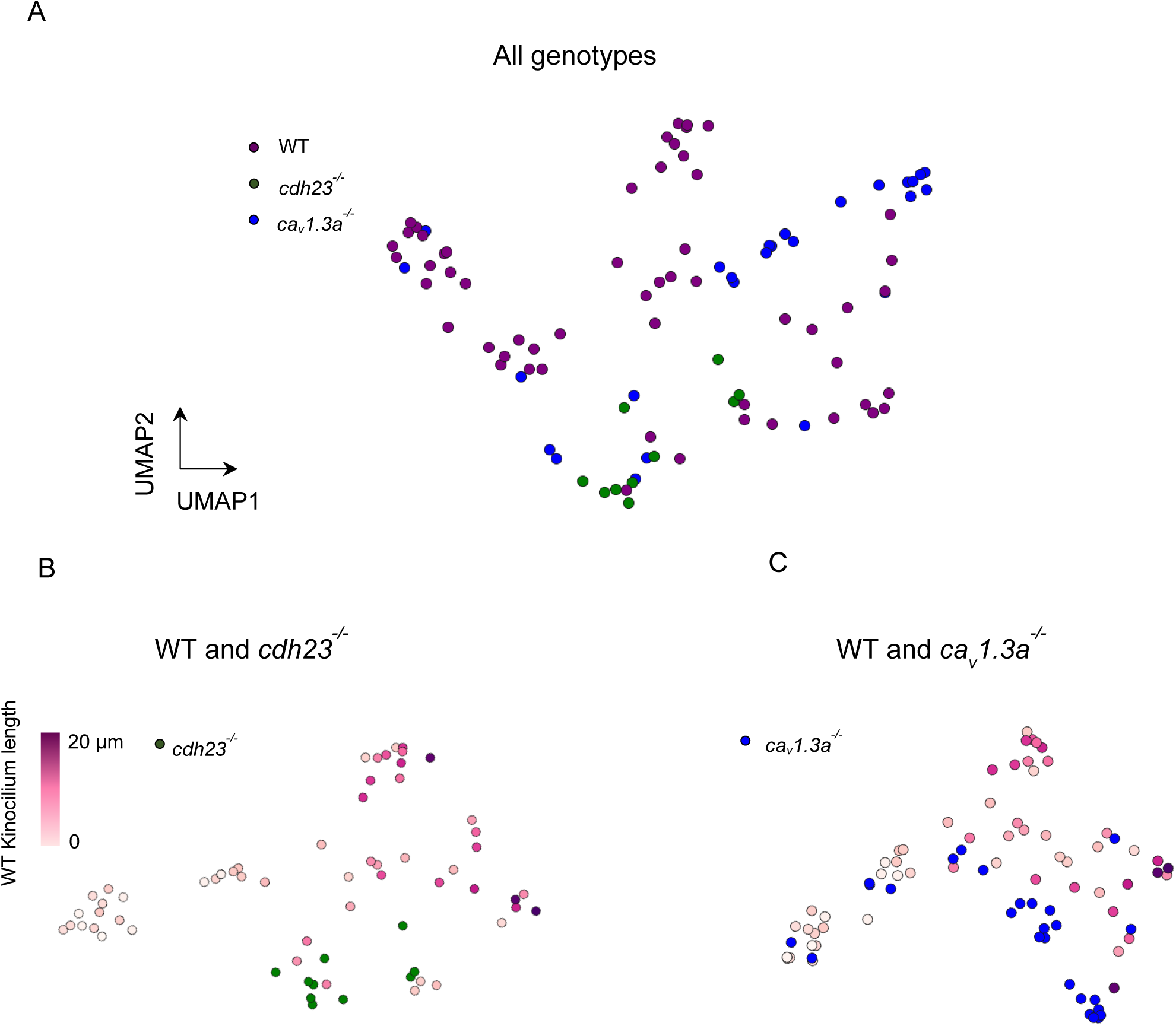
Separation of *cdh23* and *cav1.3a* hair cell mitochondrial properties from WT. (A) UMAP plot of hair cells based on principal components of their mitochondrial properties (same variables as in Figure 6). WT hair cells are in purple, *cdh23* mutants are in green, *cav1.3a* mutants are in blue. (B) UMAP analysis comparing only WT and *cdh23* hair cells. WT hair cells are color-coded by kinocilium length. (C) UMAP plot comparing WT and *cav1.3a* hair cells. WT HCs same as in Figure 6. *cdh23* HCs, n = 10, 2 NMs, 1 fish. *cav1.3a* HCs, n = 24, 2 NMs, 1 fish.

## Discussion

Although mitochondria have long been known to be essential for hair cell function, a three-dimensional ultrastructural understanding of hair cell mitochondrial architecture, and its relationship to cell function, remained understudied. We used SBFSEM to describe a hair cell-specific mitochondrial phenotype characterized by (1) high mitochondrial volume and (2) a particular mitochondrial architecture, consisting of small mitochondria apically and single large, networked mitochondria near the synaptic ribbons. There are several caveats to using SBFSEM to study mitochondrial architecture. Although it is essential for resolving individual mitochondria, SBF requires fixed tissue, providing only a snapshot of the live mitochondrial dynamics that underlie the observed architecture. In addition, the laborious nature of the reconstructions precludes multiple replicates, especially within mutant lines. However, we found no significant differences between hair cells from different neuromasts and from different WT fish (n = 5 neuromasts from 4 fish), giving us confidence that our data reflect the greater population.

We found that lateral line hair cell mitochondria contain about 40 mitochondria in total. While this is over twice the number found in the surrounding support cells, it is notably on the smaller scale compared with numbers reported for other cell types, where estimates range from hundreds to thousands (Robin and Wong, 1988; Smith and Ord, 1983). We note these estimates were made from micrographs or measurements of mtDNA, which can vary widely. There are few studies that have similar complete reconstructions of total mitochondria in cells using high resolution methodologies. A recent study indicates that there are nearly 500 mitochondria found in a primate cone photoreceptor after SBFSEM reconstruction (Hayes et al., 2021). Another recent study (Liu et al., 2022) used SBFSEM to reconstruct mitochondria from mammalian cochlear hair cells, and estimated mitochondria numbered into the thousands. A better established measurement for comparison is mitochondrial volume as a fraction of the total cell volume (Posakony et al., 1977). Our measured ratio of mitochondrial volume to total cell volume in hair cells (∼10%, Figure S1) is consistent with another recent study of mouse outer hair cells that used EM tomography and estimated mitochondrial volume about 10% of cytoplasmic volume (Perkins et al., 2020). These numbers are also in alignment with SBFSEM reconstruction of neurons and glia in rat cortex, with mitochondrial to cell volumes of 6-10% (Calì et al., 2019), though unfortunately mitochondrial number was not reported in this study.

We find distinct mitochondrial architecture along the hair cell apicobasal axis. At the hair cell apical pole, we find smaller mitochondria. In rat cochlear hair cells, apical mitochondria take up calcium during mechanotransduction (Beurg et al., 2010), contributing to robust calcium buffering mechanics to maintain cytoplasmic calcium concentrations that could otherwise affect mechanotransduction and adaptation (Ricci et al., 1998; Eatock et al., 1987). Calcium influx itself could also lead to the smaller size of mitochondria. In neurons, calcium has been reported to induce mitochondrial fission (Rintoul et al., 2003), and the fission regulator Drp1 is activated by calcium (Cribbs and Strack, 2007; Han et al., 2008; Cereghetti et al., 2008). As excess calcium uptake can lead to mitochondrial damage (Starkov et al., 2004), mitochondrial fission in this region might facilitate mitophagy and quality control (Pfluger et al., 2015; Twig et al., 2008). The small size may also provide increased efficiency in packing, helping to maintain distinct calcium pools in apical and basal compartments, as has been suggested for photoreceptor mitochondria (Giarmarco et al., 2017). Mitochondria in *opa1* mutants, which are uniformly small, demonstrated reduced calcium buffering. This could be a result of their small size, or diminished mitochondrial health from lack of fusion, preventing resource sharing and hence leading to depolarized potentials.

At the basolateral pole, hair cells contained large mitochondrial networks associated with ribbon synapses. Synaptic transmission, involving processes such as vesicle recycling and ATP-dependent removal of calcium from the cytoplasm via PMCA pumps, requires large energetic expenditures (Li and Sheng, 2022). The proximity of the networked mitochondria to the ribbons suggests these large mitochondria serve as active metabolic support. Mitochondrial fusion boosts oxidative phosphorylation and sharing of mitochondrial resources (Picard et al., 2013), indicating that the mitochondrial networks we observe would be particularly well-suited for this purpose. Synaptic mitochondria have been shown to take up calcium that enters through L-type CaV1.3 calcium channels, and disrupting this mitochondrial calcium uptake deregulated synaptic transmission (Wong et al., 2019). Larger mitochondria also have greater capacity to take up calcium (Kowaltowski et al., 2019; Szabadkai et al., 2006). As mitochondrial calcium uptake stimulates ATP production, interactions between mitochondrial fusion and synapse activity would be well-tuned to the cell’s energetic demands.

Our findings show that neither proper mechanotransduction nor synaptic activity are necessary for the growth in mitochondrial volume during hair cell maturation. This is surprising, considering the high metabolic load these processes incur. Hair cells rely on oxidative phosphorylation for 75% of their metabolic needs (Puschner and Schacht, 1997), and changes in metabolic activity is a primary driver for mitochondrial biogenesis in many tissues (Jornayvaz and Shulman, 2010). Moreover, the role of calcium influx stimulating mitochondrial biogenesis in many other electrically active cell types, such as skeletal muscle and cardiomyocytes, is well established (Ojuka et al., 2002; Chin, 2004). We note that *cdh23* mutations used to disrupt mechanotransduction still exhibit low levels of spontaneous synaptic release (Trapani and Nicolson, 2011), and cannot rule out the possibility that this residual activity might be sufficient to promote biogenesis. However, we can conclude that growth of the mitochondrial volume is not regulated by overall activity levels. Additionally, we found that disrupting the mitochondrial architecture with a mutation in *opa1* has no effect on the total mitochondrial volume. These fragmented mitochondria demonstrated depolarized potentials and decreased calcium uptake, likely associated with reduced ATP production. Together, these results suggest that the developmental increase in mitochondrial biogenesis is robust to changes in metabolic demands.

As the youngest hair cells have a total mitochondrial volume similar to that of peripheral support cells, which serve as hair cell progenitors (Thomas and Raible, 2019), we suggest that the increase in mitochondrial volume is not linked to initial cell fate specification. Supporting this, the total mitochondrial volume increases gradually as hair cells mature. This developmental mitochondrial growth is not driven by changes in cell volume, as described in other cell types (Rafelski et al., 2012; Miettinen and Björklund, 2017). Instead, it must be a product of different, ongoing pathways. One possibility may include the sirtuin deacetylases (SIRTs), key sensors of metabolism (Nogueiras et al., 2012) that can upregulate mitochondrial biogenesis through a series of parallel pathways (Yuan et al., 2016). In support of this, SIRT1 has been shown to be highly expressed in cochlear inner ear hair cells (Xiong et al., 2014), and upregulation of SIRT1 pathways is protective against various modes of hair cell death (Zhan et al., 2021; Liang et al., 2021). The steady expansion of mitochondrial volume suggests these biogenesis-promoting pathways outweigh mitophagy and quality control. Such high mitochondrial volumes may produce dangerous reactive oxygen species (ROS) levels as a byproduct of oxidative phosphorylation (Zhu et al., 2013). Therefore, the high mitochondrial volume might render hair cells vulnerable to outside stresses that would additionally increase intracellular ROS. The observed mitochondrial networks might also in part counteract this vulnerability. Promoting mitochondrial fusion is protective against starvation mediated apoptosis (Gomes et al., 2011) and SIRTs also regulate mitochondrial fusion (Uddin et al., 2021). SIRT3, located to mitochondria, has been implicated in counteracting age-related hearing loss and noise induced damage (Someya and Prolla, 2010; Brown et al., 2014; Patel et al., 2020).

We found that mechanotransduction, while having little effect on mitochondrial biogenesis, was necessary for mitochondrial architecture. The clustered position of mechanotransduction mutants about midway within the mitochondrial UMAP trajectory (Figure 10) suggests a state of incomplete development where cells are unable to progress further. Lack of mechanotransduction resulted in fewer, uniformly sized mitochondria, with reduced numbers of small apical mitochondria. This is consistent with a model where apical calcium through mechanotransduction channels promotes mitochondrial fission. Mitochondria in mechanotransduction mutants also did not form large basolateral networks localizing to synaptic ribbons. The fact that interference with mechanotransduction has effects on both formation of small mitochondria and localization of large, single networks suggests complex regulation of mitochondrial morphology.

By contrast, basal calcium entry associated with synaptic transmission is necessary for formation of basal mitochondrial networks, but not formation of smaller apical mitochondria, which were indeed present in excess numbers in *cav1.3a* mutants. In UMAP space, *cav1.3a* mutants intersperse with younger WT hair cells, but segregate from mature hair cells (Figure 10). Previous work (Trapani and Nicolson, 2011) has shown that *cav1.3a* mutants demonstrate reduced microphonic potentials reflecting decreased mechanotransduction. If calcium influx drives apical mitochondrial fission, it is present at a level enough to do so in these mutants. Basal mitochondria take up calcium during synaptic transmission (Wong et al., 2019), suggesting a paradox where calcium might promote both mitochondrial fission apically and fusion basally. However, synaptic transmission requires large energy expenditures in addition to calcium regulation, and these demands may instead drive mitochondrial fusion into basal networks. Understanding the paradoxical effects of hair cell activity influencing both the formation of smaller apical mitochondria and larger basal networks will require additional study.

We find that altering calcium entry and synaptic transmission also altered ribbon size and morphology. These results compare with previous work showing that *cav1.3a* mutant hair cells have enlarged ribbons (Sheets et al., 2012). Wong et al. (2019) demonstrated that blocking mitochondrial calcium entry also resulted in larger ribbons, a phenotype mimicked by altering NAD^+^/NADH ratios. While both of these studies showed larger ribbons after manipulating calcium and mitochondrial function, we observed *cav1.3a* mutant ribbons to be on average smaller than WT, but frequently found in clusters. We believe this discrepancy might be accounted for by the fact that the smaller ribbons we found in clusters at the EM level would appear to be larger single ribbons using fluorescence light microscopy methods employed in these previous studies. We also used strict, structural parameters to define ribbons, which might account for differences in observed ribbon numbers and their locations as seen in fluorescence (Wong et al., 2019).

Not all hair cells within zebrafish neuromasts are synaptically active, with active hair cells tending to be younger cells on the neuromast periphery (Zhang et al., 2018; Wong et al., 2019; Lukasz et al., 2022). This is interesting, considering we find the largest mitochondrial networks in the central, older hair cells. This discrepancy might be explained by considering that the initial onset of calcium entry associated with synaptic transmission might be sufficient to drive mitochondrial fusion, and that basolateral mitochondria remain networked regardless of whether the hair cell returns to a synaptically silent state during maturation.

Overall, our study provides a high resolution, three-dimensional picture of zebrafish hair cell mitochondria, and demonstrates that through mechanotransduction and synaptic activity, these cells develop a finely-tuned mitochondrial phenotype reflective of their function. This hair cell phenotype, through its high mitochondrial volume and large, networked mitochondria with efficient calcium uptake, might be necessary to support high metabolic demands, but could also increase vulnerability to small fluctuations in intracellular ROS. Disruption of this phenotype could lead to improper hair cell physiology. Thus, it is critical that future studies take into account the high specificity with which hair cells regulate their mitochondria.

## Materials and Methods

### Zebrafish

All experiments were done in compliance with the University of Washington Institutional Animal Use and Care Committee (IACUC protocol number 2997-01). Experiments were conducted when fish were 5 days post fertilization (dpf) unless otherwise noted. Sex is not determined at this age. Larvae were raised in embryo medium (14.97 mM NaCl, 500 µM KCl, 42 µM Na_2_HPO4, 150 µM KH_2_PO_4_, 1 mM CaCl_2_ dehydrate, 1 mM MgSO_4_, 0.714 mM NaHCO_3_, pH 7.2) at 28.0° C.

### Genetics lines

WT animals were of the AB strain. Both *cdh23*^tj264^ (ZDB-GENE-040413-7) and *cav1.3a*^tc323d^ (also known as *cacna1da*, ZDB-GENE-030616-135) mutant lines were generously received from the Nicolson lab (Stanford, Palo Alto, CA, USA). There is a possibility one WT fish used in SBF studies was heterozygous for *cdh23*, but this should have no impact on results (Pickett et al. 2018). Tg[*myo6b:mitoGFP*]^w213^ fish express GFP targeted to the mitochondrial matrix via a cytochrome C oxidase subunit VIII localization sequence driven by a hair cell specific promoter. Tg[*myo6b:MITO-GCaMP*]^w119^ fish express GCaMP3 targeted to the mitochondrial matrix. These fish have been characterized by our lab (Esterberg et al., 2014; Pickett et al., 2018).

The *opa1* mutant (*opa1*^w264^) was generated through CRISPR Cas9 technology. CRISPR guides were designed via http://crisperscan.org targeting *opa1* (mitochondrial dynamin like GTPase, ZDB-GENE-041114-7). Two gRNAs were prepared, with sequences GGCGAGACGGGCCACCAGA and GGCAGTGCGGTGGTCTCTGT. Both guides targeted the second exon. The gRNAs were prepared according to the protocol outlined in (Shah et al., 2015), and purified with a Zymo RNA concentrator kit. gRNAs were diluted to 1 µg/µl and stored at -80°C. Both guides were mixed with Cas9 protein (PNA Bio, Newbury Park, CA, USA) and simultaneously injected into embryos at the single-cell stage. Isolation and sequencing of a single allele showed a fifteen base pair insertion containing a premature stop codon 234 base pairs into the second exon (sequence with insert underlined, stop codon in bold: GACCTTCTG<u>T**TGA**TCGGACCTTCG</u>GGTGGCCCGTCTC), resulting in a truncated protein of 88 amino acids.

### Serial block-face scanning electron microscopy

Fish were fixed in 4% glutaraldehyde prepared with 0.1 M sodium cacodylate (pH 7.4) overnight, then decapitated. Heads were fixed for an additional night. Samples were then processed using a modified version of the Ellisman protocol (Deerinck et al. 2022). Briefly, samples were treated in 4% osmium tetroxide for 1 hr. After washout, samples were infiltrated with epon resin and baked at 60°C for 48 hrs. Transverse semithin sections were taken beginning at the anterior of the fish to localize NMs SO1 or SO2 (Raible and Kruse, 2000), above the eye. The beginning of a neuromast was identifiable by the presence of a small clustering of support cells. Samples were then mounted onto a pin and placed into the SBFSEM Volumescope (Apreo, Thermo Fisher Scientific). Images were collected with a pixel size of 5 nm. Slices were 40-50 nm thick. Typically, both the right and left corresponding neuromasts could be imaged. Hair cells, support cells, and their mitochondria were reconstructed in TrakEM2.0 (Fiji) via manual segmentation. Volume measurements were performed in AMIRA 6.5 for EM Systems (Thermo Fisher Scientific). Measures of kinocilium length were performed in TrakEM2.0. Measurements of object position within the neuromast were performed in TrakEM 2.0, and their relative geometric distances calculated using a Microsoft Excel script. No masking was used during analysis.

### Waterjet and calcium imaging

A waterjet assay was used to stimulate hair cells and record their calcium responses. Fish (5-6 dpf) were immobilized by MESAB and positioned under a harp so that neuromasts were readily accessible. Neuromasts were imaged with a 63X water immersion objective under brightfield. A glass pipette was filled with extracellular solution was placed approximately 100 µm from the neuromast. Proper positioning of the waterjet pipette was confirmed by visualizing movement of the kinocilium at the apical end of the neuromast. After a 10 sec baseline, a 10 Hz sinusoidal pressure wave was then applied using a pressure clamp (HSPC-1, ALA Scientific) for 20 sec. Images were taken with a 488 nm laser in 1s intervals using Slidebook (Intelligent Imaging Innovations). Exposure time was 100 ms. Timelapses were analyzed in Slidebook. Cells that demonstrated signal rundown during baseline were omitted from analysis.

### Confocal microscopy

TMRE dye (Invitrogen) was prepared according to manufacturer’s specifications. Tg[*myo6b:mitoGFP*]^w213^ fish (5 dpf) were allowed to swim freely in EM containing the dye (1 nM) for 1 hr prior to imaging, then immobilized with MESAB and mounted onto coverslips with 1.5% ultrapure agarose. Primary posterior lateral line neuromasts were imaged with an Zeiss 880 confocal with Airyscan technology and a 40X water immersion objective. Z-stacks were taken through the neuromasts in 0.22 µm steps. Whole neuromasts were analyzed using IMARIS software (Oxford Instruments) by creating 3D masks from the green channel to measure mean fluorescence from the red channel.

### Data analysis

Principal component analysis and two-dimensional UMAP analysis were conducted and statistically analyzed using Python 3.9.7. All other statistical analyses were conducted using GraphPad Prism 9.3.1 software (Dotmatics).

## Acknowledgements

The authors thank David White and the UW Fish Facility staff for animal care. We thank Teresa Nicolson for the *cdh23* and *cav1.3a* mutants. Lastly, the authors thank Rachel Wong for initially providing equipment and training in SBF reconstructions, and for thoughtful comments on the manuscript.

## Additional Information

### Funding

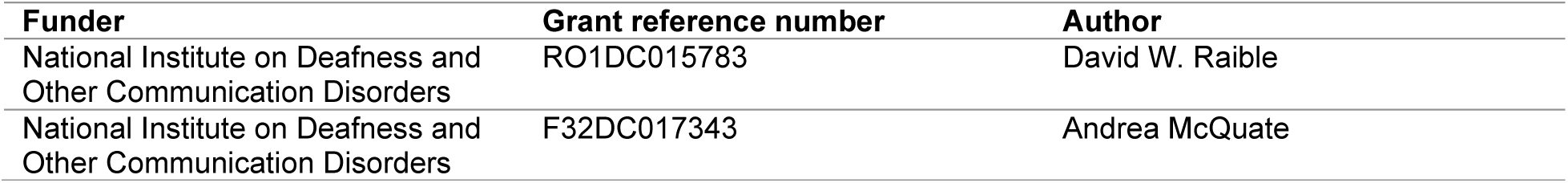

The funders had no role in study design, data collection and interpretation, or the decision to submit the work for publication

### Author contributions

Andrea McQuate: Conceptualization, data collection, formal analysis including all reconstructions, investigation, methodology, writing—original draft. Sharmon Knecht: Data collection, expertise in electron microscopy. David W. Raible: Conceptualization, formal analysis, supervision, funding acquisition, project administration, writing—review and editing.

### Competing interests

The authors declare no competing financial or non-financial interests.

### Materials/Data availability statement

The original SBFSEM data volumes for the fourteen different neuromasts used in this manuscript, including WT and mutants, will be deposited for public access at the webKnossos site (http://demo.webknossos.org/) as of July 2022.

## Figures

**Figure S1:**
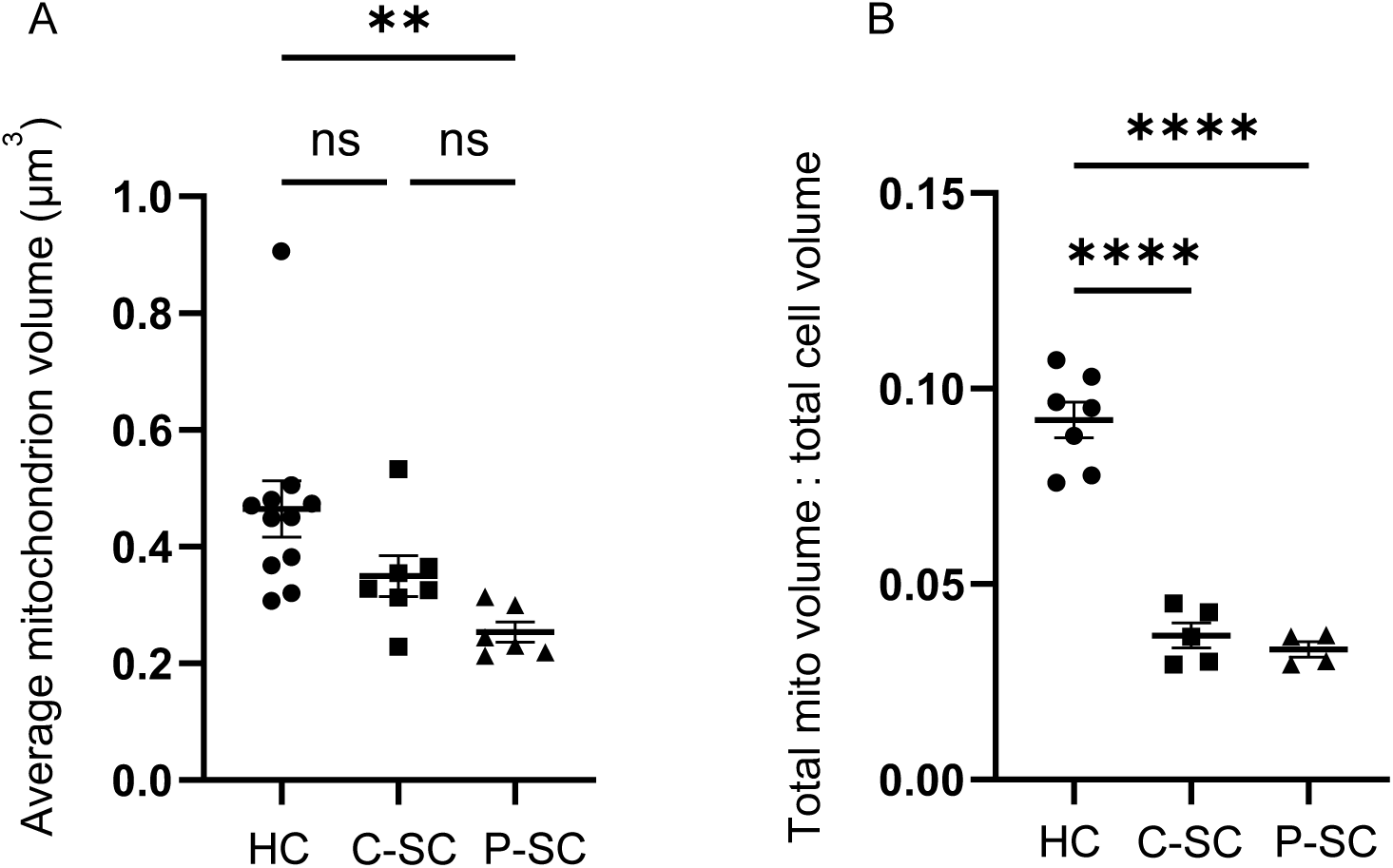
Hair cells have a higher ratio of mitochondria to cell volume, but not on average larger mitochondria, than support cells. (A) Average mitochondrion volumes in hair cells, central support cells, and peripheral support cells. (In µm^3^) HC: 0.46 ± 0.04 (n = 11, 3 NM, 3 fish), C-SC: 0.35 ± 0.03 (n = 7, 2 NM from different fish), P-SC: 0.25 ± 0.02 (n =6, 2 NM from different fish). (B) Ratio of total mitochondrial volume to cell volume in hair cells, central support cells, and peripheral support cells. Percentage of mitochondrial volume to cell volume: HC: 9.1 ± 0.4 (n =7, 2 NM from different fish, C-SC: 3.6 ± 0.3 (n = 5, 2 NM from different fish), P-SC: 3.3 ± 0.2 (n=4, 1 NM). Both: data presented as mean ± SEM, one-way ANOVA with Tukey’s multiple comparisons, ** p < 0.01, **** p < 0.0001.

**Figure S2:**
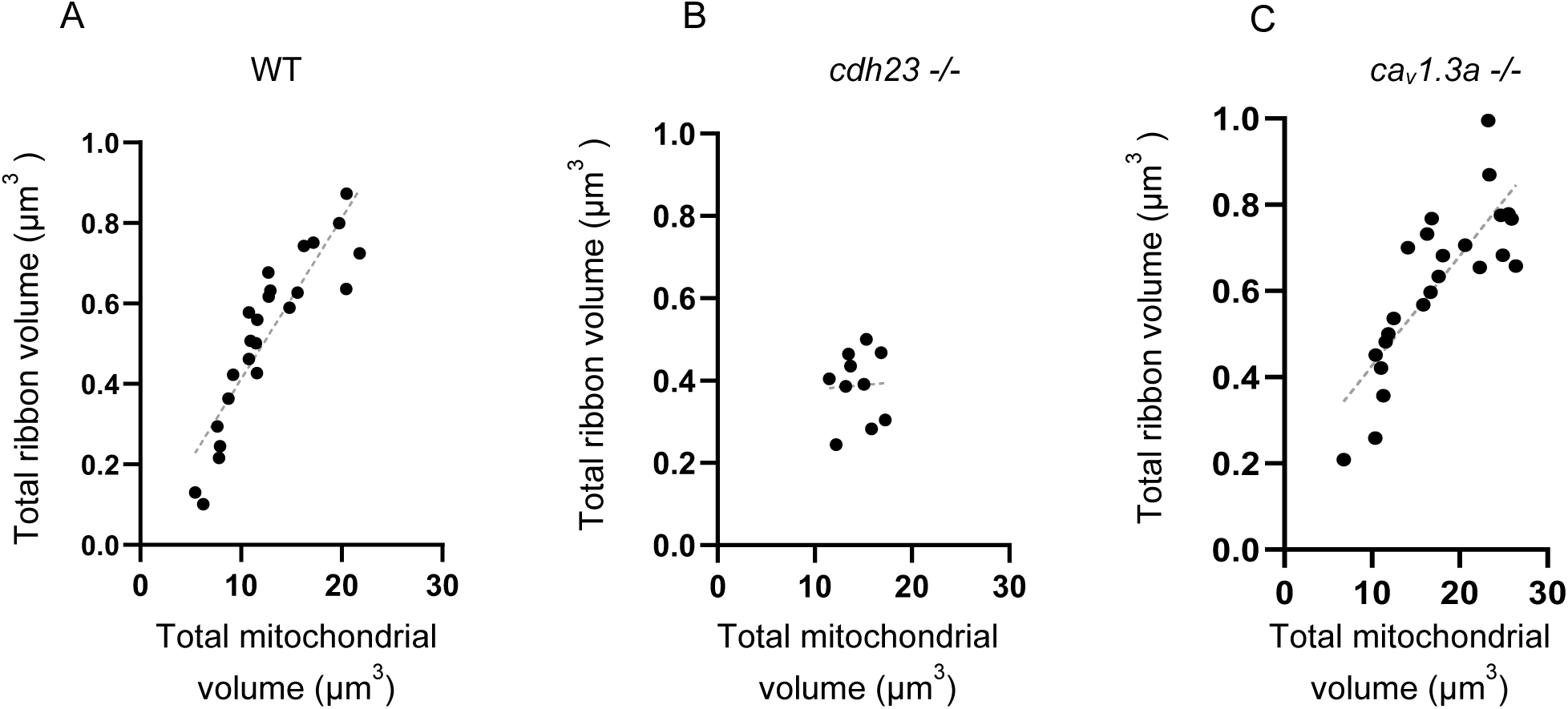
Total HC mitochondrial volume correlates with total ribbon volume. (A) Relationship between total mitochondrial volume and ribbon volume in WT (R^2^ = 0.8, p < 0.0001, n = 24, 2, 5dpf NM from different fish). (B) Relationship between total mitochondrial volume and total ribbon volume in *cdh23* mutants (R^2^ = 0.002, p = 0.9, n = 10, 2 NMs, 1 fish). (C) Relationship between total mitochondrial volume and total ribbon volume in *cav1.3a* mutants (R^2^ = 0.7, p < 0.0001, n = 24, 2, 5dpf NM, 1 fish). Gray line: linear regression.

**Figure S3:**
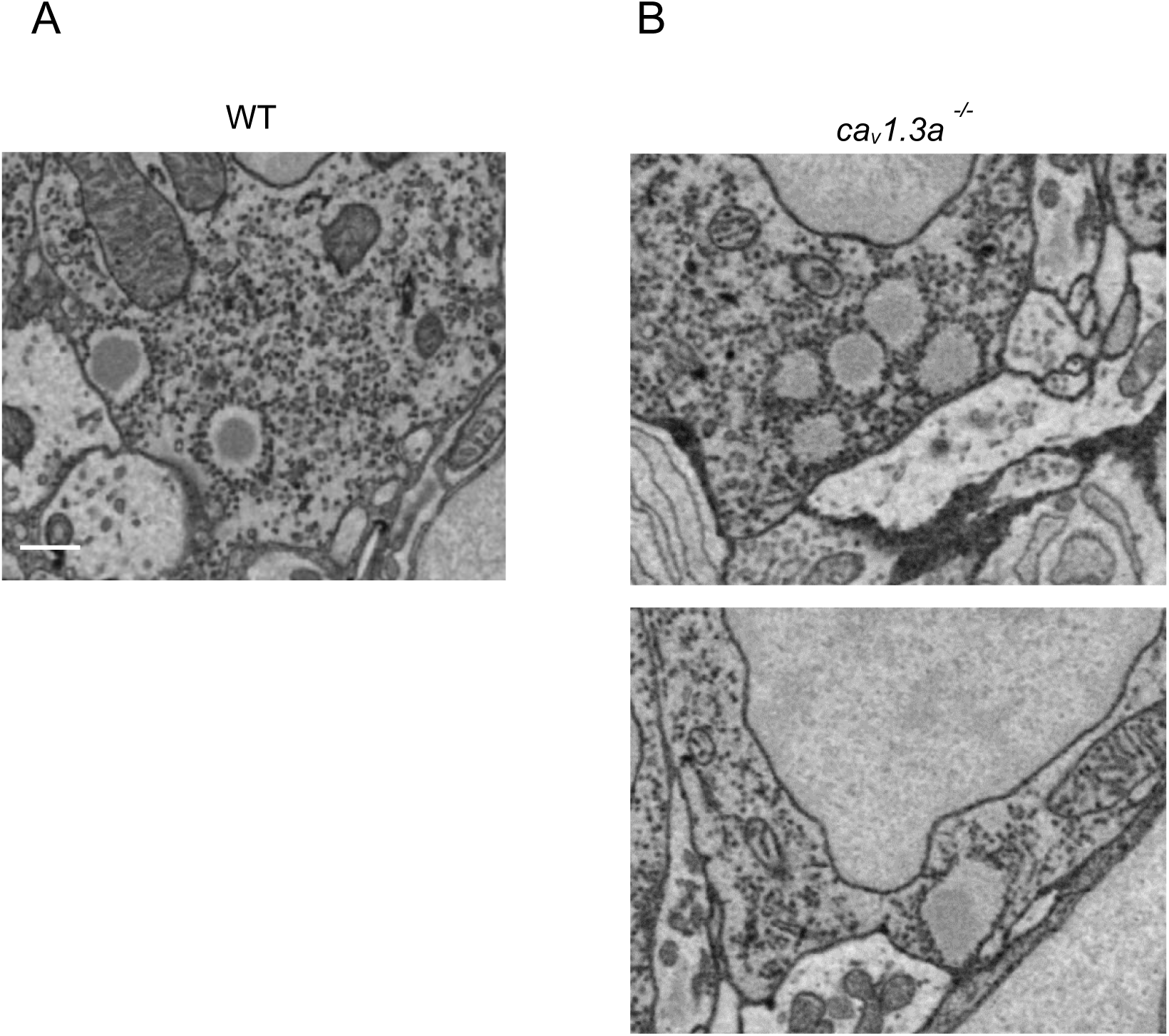
*cav1.3a* hair cells demonstrate abnormal ribbon characteristics. (A) Example of WT ribbons. (B) (Top) A cluster of ribbons from a *cav1.3a* hair cell (Bottom) polarized positioning of ribeye in a large *cav1.3a* ribbon. Scale bar: 500 nm.

## Notes

### Competing Interest Statement

The authors have declared no competing interest.

## References

Beurg, M., Nam, J.-H., Chen, Q., and Fettiplace, R. (2010). Calcium Balance and Mechanotransduction in Rat Cochlear Hair Cells. J Neurophysiol 104, 18–34. https://doi.org/10.1152/jn.00019.2010.

Böttger, E.C., and Schacht, J. (2013). The mitochondrion: A perpetrator of acquired hearing loss. Hearing Res 303, 12–19. https://doi.org/10.1016/j.heares.2013.01.006.

Brown, K.D., Maqsood, S., Huang, J.-Y., Pan, Y., Harkcom, W., Li, W., Sauve, A., Verdin, E., and Jaffrey, S.R. (2014). Activation of SIRT3 by the NAD+ Precursor Nicotinamide Riboside Protects from Noise-Induced Hearing Loss. Cell Metab 20, 1059–1068. https://doi.org/10.1016/j.cmet.2014.11.003.

Calì, C., Agus, M., Kare, K., Boges, D.J., Lehväslaiho, H., Hadwiger, M., and Magistretti, P.J. (2019). 3D cellular reconstruction of cortical glia and parenchymal morphometric analysis from Serial Block-Face Electron Microscopy of juvenile rat. Prog Neurobiol 183, 101696. https://doi.org/10.1016/j.pneurobio.2019.101696.

Cereghetti, G.M., Stangherlin, A., Brito, O.M. de, Chang, C.R., Blackstone, C., Bernardi, P., and Scorrano, L. (2008). Dephosphorylation by calcineurin regulates translocation of Drp1 to mitochondria. Proc National Acad Sci 105, 15803–15808. https://doi.org/10.1073/pnas.0808249105.

Chin, E.R. (2004). The role of calcium and calcium/calmodulin-dependent kinases in skeletal muscle plasticity and mitochondrial biogenesis. P Nutr Soc 63, 279–286. https://doi.org/10.1079/pns2004335.

Cribbs, J.T., and Strack, S. (2007). Reversible phosphorylation of Drp1 by cyclic AMP-dependent protein kinase and calcineurin regulates mitochondrial fission and cell death. Embo Rep 8, 939–944. https://doi.org/10.1038/sj.embor.7401062.

Deerinck, T.J., Bushong, E.A., Ellisman, M.H., and Thor, A. Preparation of Biological Tissues for Serial Block Face Scanning Electron Microscopy (SBEM) v2. https://doi.org/10.17504/protocols.io.36wgq7je5vk5/v2.

Eatock, R., Corey, D., and Hudspeth, A. (1987). Adaptation of mechanoelectrical transduction in hair cells of the bullfrog’s sacculus. J Neurosci 7, 2821–2836. https://doi.org/10.1523/jneurosci.07-09-02821.1987.

Esterberg, R., Hailey, D.W., Rubel, E.W., and Raible, D.W. (2014). ER–Mitochondrial Calcium Flow Underlies Vulnerability of Mechanosensory Hair Cells to Damage. J Neurosci 34, 9703–9719. https://doi.org/10.1523/jneurosci.0281-14.2014.

Esterberg, R., Linbo, T., Pickett, S.B., Wu, P., Ou, H.C., Rubel, E.W., and Raible, D.W. (2016). Mitochondrial calcium uptake underlies ROS generation during aminoglycoside-induced hair cell death. J Clin Invest 126, 3556–3566. https://doi.org/10.1172/jci84939.

Giarmarco, M.M., Cleghorn, W.M., Sloat, S.R., Hurley, J.B., and Brockerhoff, S.E. (2017). Mitochondria Maintain Distinct Ca2+ Pools in Cone Photoreceptors. J Neurosci 37, 2061–2072. https://doi.org/10.1523/jneurosci.2689-16.2017.

Gomes, L.C., Benedetto, G.D., and Scorrano, L. (2011). During autophagy mitochondria elongate, are spared from degradation and sustain cell viability. Nat Cell Biol 13, 589–598. https://doi.org/10.1038/ncb2220.

Han, X.-J., Lu, Y.-F., Li, S.-A., Kaitsuka, T., Sato, Y., Tomizawa, K., Nairn, A.C., Takei, K., Matsui, H., and Matsushita, M. (2008). CaM kinase Iα–induced phosphorylation of Drp1 regulates mitochondrial morphology. J Cell Biology 182, 573–585. https://doi.org/10.1083/jcb.200802164.

Hayes, M.J., Tracey-White, D., Kam, J.H., Powner, M.B., and Jeffery, G. (2021). The 3D organisation of mitochondria in primate photoreceptors. Sci Rep-Uk 11, 18863. https://doi.org/10.1038/s41598-021-98409-7.

Jornayvaz, F.R., and Shulman, G.I. (2010). Regulation of mitochondrial biogenesis. Essays Biochem 47, 69–84. https://doi.org/10.1042/bse0470069.

Kindt, K.S., Finch, G., and Nicolson, T. (2012). Kinocilia Mediate Mechanosensitivity in Developing Zebrafish Hair Cells. Dev Cell 23, 329–341. https://doi.org/10.1016/j.devcel.2012.05.022.

Kokotas, H., Petersen, M., and Willems, P. (2007). Mitochondrial deafness. Clin Genet 71, 379–391. https://doi.org/10.1111/j.1399-0004.2007.00800.x.

Kowaltowski, A.J., Menezes-Filho, S.L., Assali, E.A., Gonçalves, I.G., Cabral-Costa, J.V., Abreu, P., Miller, N., Nolasco, P., Laurindo, F.R.M., Bruni-Cardoso, A., et al. (2019). Mitochondrial morphology regulates organellar Ca 2+ uptake and changes cellular Ca 2+ homeostasis. Faseb J 33, 13176–13188. https://doi.org/10.1096/fj.201901136r.

Lesus, J., Arias, K., Kulaga, J., Sobkiv, S., Patel, A., Babu, V., Kambalyal, A., Patel, M., Padron, F., Mozaffari, P., et al. (2019). Why study inner ear hair cell mitochondria? Hno 67, 429–433. https://doi.org/10.1007/s00106-019-0662-2.

Li, S., and Sheng, Z.-H. (2022). Energy matters: presynaptic metabolism and the maintenance of synaptic transmission. Nat Rev Neurosci 23, 4–22. https://doi.org/10.1038/s41583-021-00535-8.

Liang, Z., Zhang, T., Zhan, T., Cheng, G., Zhang, W., Jia, H., and Yang, H. (2021). Metformin alleviates cisplatin-induced ototoxicity by autophagy induction possibly via the AMPK/FOXO3a pathway. J Neurophysiol 125, 1202–1212. https://doi.org/10.1152/jn.00417.2020.

Liu, J., Wang, S., Lu, Y., Wang, H., Wang, F., Qiu, M., Xie, Q., Han, H., and Hua, Y. (2022). Aligned Organization of Synapses and Mitochondria in Auditory Hair Cells. Neurosci Bull 38, 235–248. https://doi.org/10.1007/s12264-021-00801-w.

Liu, Y.J., McIntyre, R.L., Janssens, G.E., and Houtkooper, R.H. (2020). Mitochondrial fission and fusion: A dynamic role in aging and potential target for age-related disease. Mech Ageing Dev 186, 111212. https://doi.org/10.1016/j.mad.2020.111212.

Lukasz, D., Beirl, A., and Kindt, K. (2022). Chronic neurotransmission increases the susceptibility of lateral-line hair cells to ototoxic insults. Biorxiv 2022.02.22.481465. https://doi.org/10.1101/2022.02.22.481465.

Miettinen, T.P., and Björklund, M. (2017). Mitochondrial Function and Cell Size: An Allometric Relationship. Trends Cell Biol 27, 393–402. https://doi.org/10.1016/j.tcb.2017.02.006.

Nogueiras, R., Habegger, K.M., Chaudhary, N., Finan, B., Banks, A.S., Dietrich, M.O., Horvath, T.L., Sinclair, D.A., Pfluger, P.T., and Tschöp, M.H. (2012). Sirtuin 1 and Sirtuin 3: Physiological Modulators of Metabolism. Physiol Rev 92, 1479–1514. https://doi.org/10.1152/physrev.00022.2011.

Ojuka, E.O., Jones, T.E., Han, D.-H., Chen, M., Wamhoff, B.R., Sturek, M., and Holloszy, J.O. (2002). Intermittent increases in cytosolic Ca2+stimulate mitochondrial biogenesis in muscle cells. Am J Physiol-Endoc M 283, E1040–E1045. https://doi.org/10.1152/ajpendo.00242.2002.

Owens, K.N., Cunningham, D.E., Macdonald, G., Rubel, E.W., Raible, D.W., and Pujol, R. (2007). Ultrastructural analysis of aminoglycoside-induced hair cell death in the zebrafish lateral line reveals an early mitochondrial response. J Comp Neurol 502, 522–543. https://doi.org/10.1002/cne.21345.

Patel, S., Shah, L., Dang, N., Tan, X., Almudevar, A., and White, P.M. (2020). SIRT3 promotes auditory function in young adult FVB/nJ mice but is dispensable for hearing recovery after noise exposure. Plos One 15, e0235491. https://doi.org/10.1371/journal.pone.0235491.

Perkins, G., Lee, J.H., Park, S., Kang, M., Perez-Flores, M.C., Ju, S., Phillips, G., Lysakowski, A., Gratton, M.A., and Yamoah, E.N. (2020). Altered Outer Hair Cell Mitochondrial and Subsurface Cisternae Connectomics Are Candidate Mechanisms for Hearing Loss in Mice. J Neurosci 40, 8556–8572. https://doi.org/10.1523/jneurosci.2901-19.2020.

Perkins, G.A., Tjong, J., Brown, J.M., Poquiz, P.H., Scott, R.T., Kolson, D.R., Ellisman, M.H., and Spirou, G.A. (2010). The Micro-Architecture of Mitochondria at Active Zones: Electron Tomography Reveals Novel Anchoring Scaffolds and Cristae Structured for High-Rate Metabolism. J Neurosci 30, 1015–1026. https://doi.org/10.1523/jneurosci.1517-09.2010.

Pfluger, P.T., Kabra, D.G., Aichler, M., Schriever, S.C., Pfuhlmann, K., García, V.C., Lehti, M., Weber, J., Kutschke, M., Rozman, J., et al. (2015). Calcineurin Links Mitochondrial Elongation with Energy Metabolism. Cell Metab 22, 838–850. https://doi.org/10.1016/j.cmet.2015.08.022.

Picard, M., Shirihai, O.S., Gentil, B.J., and Burelle, Y. (2013). Mitochondrial morphology transitions and functions: implications for retrograde signaling? Am J Physiology-Regulatory Integr Comp Physiology 304, R393–R406. https://doi.org/10.1152/ajpregu.00584.2012.

Pickett, S.B., and Raible, D.W. (2019). Water Waves to Sound Waves: Using Zebrafish to Explore Hair Cell Biology. J Assoc Res Otolaryngology 20, 1–19. https://doi.org/10.1007/s10162-018-00711-1.

Pickett, S.B., Thomas, E.D., Sebe, J.Y., Linbo, T., Esterberg, R., Hailey, D.W., and Raible, D.W. (2018). Cumulative mitochondrial activity correlates with ototoxin susceptibility in zebrafish mechanosensory hair cells. Elife 7, e38062. https://doi.org/10.7554/elife.38062.

Posakony, J., England, J., and Attardi, G. (1977). Mitochondrial growth and division during the cell cycle in HeLa cells. J Cell Biology 74, 468–491. https://doi.org/10.1083/jcb.74.2.468.

Puschner, B., and Schacht, J. (1997). Energy metabolism in cochlear outer hair cells in vitro. Hearing Res 114, 102–106. https://doi.org/10.1016/s0378-5955(97)00163-9.

Rafelski, S.M., Viana, M.P., Zhang, Y., Chan, Y.-H.M., Thorn, K.S., Yam, P., Fung, J.C., Li, H., Costa, L. da F., and Marshall, W.F. (2012). Mitochondrial Network Size Scaling in Budding Yeast. Science 338, 822–824. https://doi.org/10.1126/science.1225720.

Raible, D.W., and Kruse, G.J. (2000). Organization of the lateral line system in embryonic zebrafish. J Comp Neurol 421, 189–198. https://doi.org/10.1002/(sici)1096-9861(20000529)421:2<189::aid-cne5>3.0.co;2-k.

Reddy, P.H., Reddy, T.P., Manczak, M., Calkins, M.J., Shirendeb, U., and Mao, P. (2011). Dynamin-related protein 1 and mitochondrial fragmentation in neurodegenerative diseases. Brain Res Rev 67, 103–118. https://doi.org/10.1016/j.brainresrev.2010.11.004.

Ricci, A.J., Wu, Y.-C., and Fettiplace, R. (1998). The Endogenous Calcium Buffer and the Time Course of Transducer Adaptation in Auditory Hair Cells. J Neurosci 18, 8261–8277. https://doi.org/10.1523/jneurosci.18-20-08261.1998.

Rintoul, G.L., Filiano, A.J., Brocard, J.B., Kress, G.J., and Reynolds, I.J. (2003). Glutamate Decreases Mitochondrial Size and Movement in Primary Forebrain Neurons. J Neurosci 23, 7881–7888. https://doi.org/10.1523/jneurosci.23-21-07881.2003.

Robin, E.D., and Wong, R. (1988). Mitochondrial DNA molecules and virtual number of mitochondria per cell in mammalian cells. J Cell Physiol 136, 507–513. https://doi.org/10.1002/jcp.1041360316.

Romero-Carvajal, A., Navajas Acedo, J., Jiang, L., Kozlovskaja-Gumbrienė, A., Alexander, R., Li, H., and Piotrowski, T. (2015). Regeneration of Sensory Hair Cells Requires Localized Interactions between the Notch and Wnt Pathways. Dev Cell 34, 267–282. https://doi.org/10.1016/j.devcel.2015.05.025.

Shah, A.N., Davey, C.F., Whitebirch, A.C., Miller, A.C., and Moens, C.B. (2015). Rapid reverse genetic screening using CRISPR in zebrafish. Nat Methods 12, 535–540. https://doi.org/10.1038/nmeth.3360.

Sheets, L., Kindt, K.S., and Nicolson, T. (2012). Presynaptic CaV1.3 Channels Regulate Synaptic Ribbon Size and Are Required for Synaptic Maintenance in Sensory Hair Cells. J Neurosci 32, 17273–17286. https://doi.org/10.1523/jneurosci.3005-12.2012.

Smith, R.A., and Ord, M.J. (1983). Mitochondrial Form and Function Relationships in Vivo: Their Potential in Toxicology and Pathology. Int Rev Cytol 83, 63–134. https://doi.org/10.1016/s0074-7696(08)61686-1.

Söllner, C., Rauch, G.-J., Siemens, J., Geisler, R., Schuster, S.C., Müller, U., and Nicolson, T. (2004). Mutations in cadherin 23 affect tip links in zebrafish sensory hair cells. Nature 428, 955–959. https://doi.org/10.1038/nature02484.

Someya, S., and Prolla, T.A. (2010). Mitochondrial oxidative damage and apoptosis in age-related hearing loss. Mech Ageing Dev 131, 480–486. https://doi.org/10.1016/j.mad.2010.04.006.

Starkov, A.A., Chinopoulos, C., and Fiskum, G. (2004). Mitochondrial calcium and oxidative stress as mediators of ischemic brain injury. Cell Calcium 36, 257–264. https://doi.org/10.1016/j.ceca.2004.02.012.

Stowers, R.S., Megeath, L.J., Górska-Andrzejak, J., Meinertzhagen, I.A., and Schwarz, T.L. (2002). Axonal Transport of Mitochondria to Synapses Depends on Milton, a Novel Drosophila Protein. Neuron 36, 1063–1077. https://doi.org/10.1016/s0896-6273(02)01094-2.

Szabadkai, G., Simoni, A.M., Bianchi, K., Stefani, D.D., Leo, S., Wieckowski, M.R., and Rizzuto, R. (2006). Mitochondrial dynamics and Ca2+ signaling. Biochimica Et Biophysica Acta Bba - Mol Cell Res 1763, 442–449. https://doi.org/10.1016/j.bbamcr.2006.04.002.

Thomas, E.D., and Raible, D.W. (2019). Distinct progenitor populations mediate regeneration in the zebrafish lateral line. Elife 8, e43736. https://doi.org/10.7554/elife.43736.

Trapani, J.G., and Nicolson, T. (2011). Mechanism of Spontaneous Activity in Afferent Neurons of the Zebrafish Lateral-Line Organ. J Neurosci 31, 1614–1623. https://doi.org/10.1523/jneurosci.3369-10.2011.

Twig, G., Hyde, B., and Shirihai, O.S. (2008). Mitochondrial fusion, fission and autophagy as a quality control axis: The bioenergetic view. Biochimica Et Biophysica Acta Bba - Bioenergetics 1777, 1092–1097. https://doi.org/10.1016/j.bbabio.2008.05.001.

Uddin, G.M., Abbas, R., and Shutt, T.E. (2021). The role of protein acetylation in regulating mitochondrial fusion and fission. Biochem Soc T 49, 2807–2819. https://doi.org/10.1042/bst20210798.

Wang, X., Zhu, Y., Long, H., Pan, S., Xiong, H., Fang, Q., Hill, K., Lai, R., Yuan, H., and Sha, S.-H. (2019). Mitochondrial Calcium Transporters Mediate Sensitivity to Noise-Induced Losses of Hair Cells and Cochlear Synapses. Front Mol Neurosci 11, 469. https://doi.org/10.3389/fnmol.2018.00469.

Wong, H.C., Zhang, Q., Beirl, A.J., Petralia, R.S., Wang, Y.-X., and Kindt, K. (2019). Synaptic mitochondria regulate hair-cell synapse size and function. Elife 8, e48914. https://doi.org/10.7554/elife.48914.

Xiong, H., Dai, M., Ou, Y., Pang, J., Yang, H., Huang, Q., Chen, S., Zhang, Z., Xu, Y., Cai, Y., et al. (2014). SIRT1 expression in the cochlea and auditory cortex of a mouse model of age-related hearing loss. Exp Gerontol 51, 8–14. https://doi.org/10.1016/j.exger.2013.12.006.

Yuan, Y., Cruzat, V.F., Newshome, P., Cheng, J., Chen, Y., and Lu, Y. (2016). Regulation of SIRT1 in aging: Roles in mitochondrial function and biogenesis. Mech Ageing Dev 155, 10–21. https://doi.org/10.1016/j.mad.2016.02.003.

Zhan, T., Xiong, H., Pang, J., Zhang, W., Ye, Y., Liang, Z., Huang, X., He, F., Jian, B., He, W., et al. (2021). Modulation of NAD+ biosynthesis activates SIRT1 and resists cisplatin-induced ototoxicity. Toxicol Lett 349, 115–123. https://doi.org/10.1016/j.toxlet.2021.05.013.

Zhang, Q., Li, S., Wong, H.-T.C., He, X.J., Beirl, A., Petralia, R.S., Wang, Y.-X., and Kindt, K.S. (2018). Synaptically silent sensory hair cells in zebrafish are recruited after damage. Nat Commun 9, 1388. https://doi.org/10.1038/s41467-018-03806-8.

Zhu, J., Wang, K.Z., and Chu, C.T. (2013). After the banquet. Autophagy 9, 1663–1676. https://doi.org/10.4161/auto.24135.

